# A conserved enzyme of smut fungi facilitates cell-to-cell movement in the plant bundle sheath

**DOI:** 10.1101/2022.05.17.492292

**Authors:** Bilal Ökmen, Elaine Jaeger, Lena Schilling, Natalie Finke, Yoon Joo Lee, Raphael Wemhöner, Markus Pauly, Ulla Neumann, Gunther Doehlemann

**Author notes:** **Corresponding authors:** Gunther Doehlemann, Bilal Ökmen.

## Abstract

The smut fungi are one of the largest groups of fungal plant pathogens, causing disease in all cereal crops. They directly penetrate their hosts and establish a biotrophic interaction. During colonization of the plant, smut fungi secrete a wide range of effector proteins, which suppress plant immunity and modulate cellular functions as well as development of the host, thereby determining the pathogen’s life-style and virulence potential.

The conserved effector Erc1 (**<u>e</u>**nzyme **<u>r</u>**equired for **<u>c</u>**ell-to-cell movement) contributes to virulence of the corn smut *Ustilago maydis* in maize leaves, but not on the tassel. Erc1 binds to host cell wall components and has a 1,3-β-glucanase activity, which is required to attenuate β-glucan-induced defense responses in host leaves. Confocal microscopy revealed that Erc1 has a cell type-specific virulence function, being necessary for fungal cell-to-cell movement in the plant bundle sheath. This cell type-specific virulence function of Erc1 is fully conserved in the barley pathogen *Ustilago hordei*, which has a functionally conserved Erc1 orthologue.

Thus, Erc1 is an enzymatically active core virulence factor with a cell type-specific virulence function in different hosts, which is important for cell-to-cell movement during host colonization of pathogenic smut fungi.

## Introduction

Plants have established physical and chemical defense layers to protect themselves from continuous pathogenic attacks. The plant cell wall is one of the main physical barriers, which protects plants against such attacks. In turn, phytopathogenic microorganisms developed strategies to breach the host cell wall allowing successful host penetration and colonization. In this regard, some filamentous phytopathogens use both mechanical force and enzymatic activities to breach this first layer of defense. While a specialized dome-like structure, the appressorium, provides mechanical force for direct penetration, each pathogen has an arsenal of plant cell wall degrading enzymes (PCWDEs) that break down the host cell wall for host penetration or nutrient acquisition (Bradley *et al*., 2022). Plant pathogens show a wide range of variation in their PCWDEs repertoire including glycoside hydrolases (GHs), which determines their virulence, pathogenic life-styles and host specificity (Ospina-Giraldo *et al*., 2010; Zhao *et al*., 2013; Hane *et al*., 2019). Compared to necrotrophic and hemibiotrophic pathogens, biotrophs possess relatively few PCWDEs, which is in line with their need to not disturb the cell wall integrity of their host plants and minimize release of damage-associated molecular patterns (DAMPs) (de Wit *et al*., 2012). DAMPs are endogenous molecules, including plant cell wall components and peptides, that are released from a host plant upon damage or infection. They serve as signaling molecules to induce defense-related responses against invading pathogens and promote damage repair (Hou *et al*., 2019).

*Ustilago maydis*, the causative agent of corn smut disease, is a biotrophic fungal pathogen that induces tumor formation in all aerial plant organs, including leaves, ears, and tassels. Like other phytopathogenic smut fungi, it grows both extra- and intracellularly and moves from cell-to-cell to colonize its host (Zuo *et al*., 2019). To establish disease, *U. maydis* secretes effector proteins, which are deployed in an organ-specific manner (Skibbe *et al*., 2010; Schilling *et al*., 2014). Different arrangement in tissue types, different cell types, or differences in cell wall components and metabolite compositions may lead to an organ-specific function of some effectors (Matei *et al*., 2018).

Schilling *et al*., (2014) described a set of leaf specific *U. maydis* effectors of which one (UMAG_01829) is predicted to be an α-L-arabinofuranosidase, which shows the highest expression level among the *U. maydis* effector genes (Schilling *et al*., 2014; Lanver *et al*., 2018). The presence of a putative PCWDE among the organ-specific effectors might reflect differences in cell wall composition of leaves compared to tassel tissues. Many previously published reports demonstrate the importance of pathogen-derived PCWDEs in plant microbe interactions (Bradley *et al*., 2022). While some glycoside hydrolases are involved in host penetration and nutrient acquisition by degrading plant cell wall components, others are involved in detoxification of antimicrobial secondary metabolites or hydrolyzing MAMP molecules (Bradley *et al*., 2022). For example, xylanases have been shown recently to be involved in proliferation of *U. maydis* during plant infection (Moreno-Sánchez *et al*., 2021). While a GH10 family member tomatinase enzyme from *Cladosporium fulvum* is required for detoxification of tomato-specific α-tomatine (Ökmen *et al*., 2013), a GH18 family member chitinase from *Magnaporthe oryzae* (MoChia1) binds and sequesters chitin fragments released from the fungal cell wall to prevent MAMP-triggered immunity (Yang *et al*., 2019). Likewise, several arabinofuranosidases have been described to contribute to the virulence of plant pathogens. For example, the endo-arabinase BcAra1 is necessary for the full virulence of the necrotrophic plant pathogen *Botrytis cinerea* in *A. thaliana* (Nafisi *et al*., 2014). In *Sclerotinia trifoliorum*, arabinofuranosidase-deletion mutants display reduced virulence in *Pisum sativum var. avense* (Rehnstrom *et al*., 1994). α-L-arabinofuranosidases are enzymes that catalyze the hydrolysis of terminal, non-reducing α-L-arabinofuranose residues in α-L-arabinosides. The enzyme works on terminal α-L-1,2-; α-L-1,3- and α-L-1,5-arabinofuranosyl residues present in the cell wall polymers, such as arabinoxylans and arabinogalactans (Margolles-Clark *et al*., 1996). Arabinofuranosidases often act in concert with other hemicellulases to degrade the hemicellulose of the plant cell wall (Gilead & Shoham, 1995). In this study, we have functionally characterized Erc1 (**<u>e</u>**nzyme **<u>r</u>**equired for **<u>c</u>**ell-to-cell movement), a conserved effector of smut fungi with an organ-specific virulence function. Despite a predicted α-L-arabinofuranosidase activity based on protein sequence, we found that Erc1 exhibits 1,3-β-glucanase activity and is required for cell-to-cell movement specifically in bundle sheaths cells. Strikingly, this cell-type specific virulence function is conserved in both the *U. maydis*-maize and *Ustilago hordei*-barley pathosystem. This identifies Erc1 as a secreted enzyme with a highly specific virulence function, which is conserved amongst pathosystems in different plant hosts.

## Results

### UMAG_01829 is a leaf-specific virulence factor of *Ustilago maydis*

In a previous study (Schilling *et al*., 2014), we have shown organ-specific virulence activities of *U. maydis* effector genes. Within the effector genes being specifically required for tumorigenesis in maize leaves, *UMAG_01829* caught our attention because of its predicted protein properties and function, which are rather untypical for a fungal effector: *UMAG_01829* encodes a 76.4 kDa protein with 703 amino acids (aa), with an N-terminal secretion signal (1-18 aa; SignalP 5.0), followed by a predicted carbohydrate binding module (CBM, aa 124-266) **(Figure S1a)**. The C-terminal part of the protein (aa 272-691) is predicted to represent a catalytic domain of a GH Family 51 hydrolase (α-L-arabinofuranosidase) with two predicted active sites at amino acids 410-412^GNE^ and 499^E^ **(Fig. S1a**). GH family 51 hydrolases have been shown to exhibit mainly hemicellulose α-L-arabinofuranosidase activity, i.e. removing terminal arabinosyl-moieties from arabinoxylans or other branched arabinans, but also endoglucanase and endoxylanase activities have been demonstrated (cazy.org). The *U. maydis* genome has additional homologs of α-L-arabinofuranosidase (*Afu*) genes, which belong to GH51 (Afu2; UMAG_00837) and GH62 (Afu3; UMAG_04309, a commercially available α-L-arabinofuranosidase). Neither Afu2 nor Afu3 contain a predicted CBM **(Fig. S1b)** and they do not contribute to *U. maydis* virulence on maize **(Fig. S2b)** (Lanver *et al*., 2014). All three hydrolase genes are strictly expressed only *in planta* and are not expressed *in axenic* culture **(Fig. S2a)**. While *UMAG_01829* is one of the most highly expressed effector genes in *U. maydis* at all tested time points of colonization, *Afu2* is lowly expressed only at the very early and very late stage of infection (12 days post infection (dpi)) (Lanver *et al*., 2018). Like *UMAG_01829, Afu3* is expressed throughout *U. maydis* leaf infection; however, the expression level of *UMAG_01829* is significantly higher when compared to *Afu3* **(Fig. S2a)** (Lanver *et al*., 2018). Homology search and phylogenic tree analysis showed that Afu2 and Afu3 are widely conserved in fungal and bacterial genomes. In contrast, UMAG_01829 is highly conserved within Ustilaginomycotina, but it is not found outside the Basidiomycetes (**Fig. S3)**.

### *UMAG_01829* encodes an enzyme which is required for fungal cell-to-cell movement in the host bundle sheath

As a first step of functional characterization, we confirmed the previous finding of Schilling *et al*., (2014), showing that UMAG_01829 is required for full virulence of *U. maydis* in maize leaves **(Fig. 1a)**. Genetic complementation of the *U. maydis* SG200ΔUMAG_01829 mutant with the native *UMAG_01829* gene fully restored *U. maydis* virulence, confirming that the observed defect in virulence was solely caused by the deletion of *UMAG_01829*. To investigate, which step of host infection is affected by *UMAG_01829* deletion, fungal growth inside the leaf was followed by confocal microscopy. This revealed that the SG200ΔUMAG_01829 mutant displays a defect in cell-to-cell movement and this phenotype specifically appeared in bundle sheath cells **(Fig. 1b-c)**. While 67% of cell-to-cell movement attempts of the SG200ΔUMAG_01829 mutant failed in bundle sheaths cells, this was only observed in 17% for both SG200 and SG200Δerc1/C complementation strains **(Fig. 1b)**. Because of this specific function in cell-to-cell movement, we named UMAG_01829 UmErc1 (*Ustilago maydis* **<u>e</u>**nzyme **<u>r</u>**equired for **<u>c</u>**ell-to-cell movement).

**Fig. 1.**
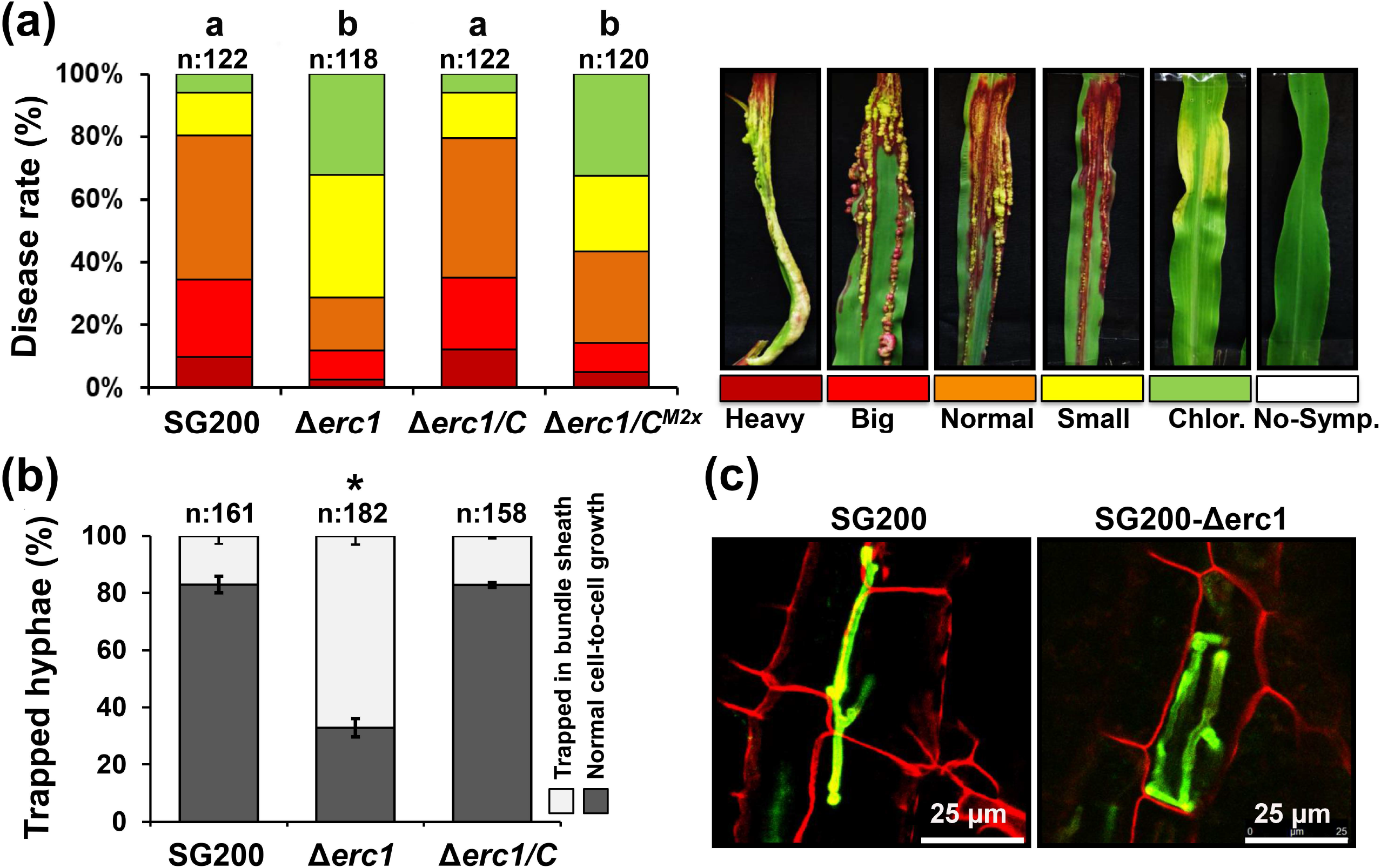
Erc1 is a virulence factor involved in cell-to-cell movement in maize. **(a)** Disease symptoms caused by *Ustilago maydis* SG200, SG200Δerc1 mutant, SG200Δerc1/C strain and SG200Δerc1/Erc1^M2x^ on EGB maize leaves at 12 days post inoculation (dpi). Disease rates are given as a percentage of the total number of infected plants. Three biological replicates were carried out. n: number of infected maize seedlings and asterisks above bars indicate significant differences (One-way ANOVA followed by Tukey multiple comparison test was performed, p<0.05). **(b)** Quantification of cell-to-cell penetration efficiency of SG200, SG200Δerc1, and SG200Δerc1/C complementation strains. The graph depicts the total number of trapped *U. maydis* hyphae in maize bundle sheath cells at 4 dpi. Three biological replicates were carried out. n: number of penetrated maize cells. Asterisks above bars indicate significant differences (p<0.05, student t-test). **(c)** Microscopic observation of trapped *U. maydis* SG200 and SG200Δerc1 hyphae in maize bundle sheath cells at 4 dpi via WGA-AF488/Propidium iodide staining. WGA-AF488 (green color -fungal cell wall): excitation at 488 nm and detection at 500-540 nm. PI (red color - plant cell wall): excitation at 561 nm and detection at 580–630 nm.

To test whether the predicted carbohydrate binding module (CBM) and its putative catalytic activity are required for its virulence function, various mutated forms of Erc1 including one without the CBM domain, as well as an active site mutant (410-412^GNE>AAA^ and 499^E>A^) were expressed in the *U. maydis* Δ*erc1* mutant background. To test for virulence complementation, maize seedlings were inoculated with *U. maydis* SG200, SG200Δerc1 mutant (Δerc1), SG200Δerc1/Erc1 (Δerc1/C), SG200Δerc1/Erc1^ΔCBM^ (ΔCBM) and SG200Δerc1/Erc1^M2X^ (Δerc1/C^M2x^) strains **(Fig. 1a and Fig. 2a)**. While complementation of wild-type Erc1 again fully recovered the virulence phenotype of SG200Δerc1 mutant at 12 dpi, the Erc1^M2X^ and Erc1^ΔCBM^ complementation only partially rescued the reduced virulence phenotype of the SG200Δerc1 mutant at 12 dpi **(Fig. 1a and Fig. 2a)**. Site-directed mutagenesis leading to amino acid exchanges at two predicted active sites of the catalytic domain (410-412^GNE>AAA^ and 499^E>A^) resulted in a significant reduction in virulence compared to the progenitor strain. However, there was no significant difference between the SG200Δerc1 and SG200Δerc1/Erc1^M2X^ strains **(Fig. 1a)**, indicating that enzymatic activity of Erc1 is required for its virulence function.

**Fig. 2.**
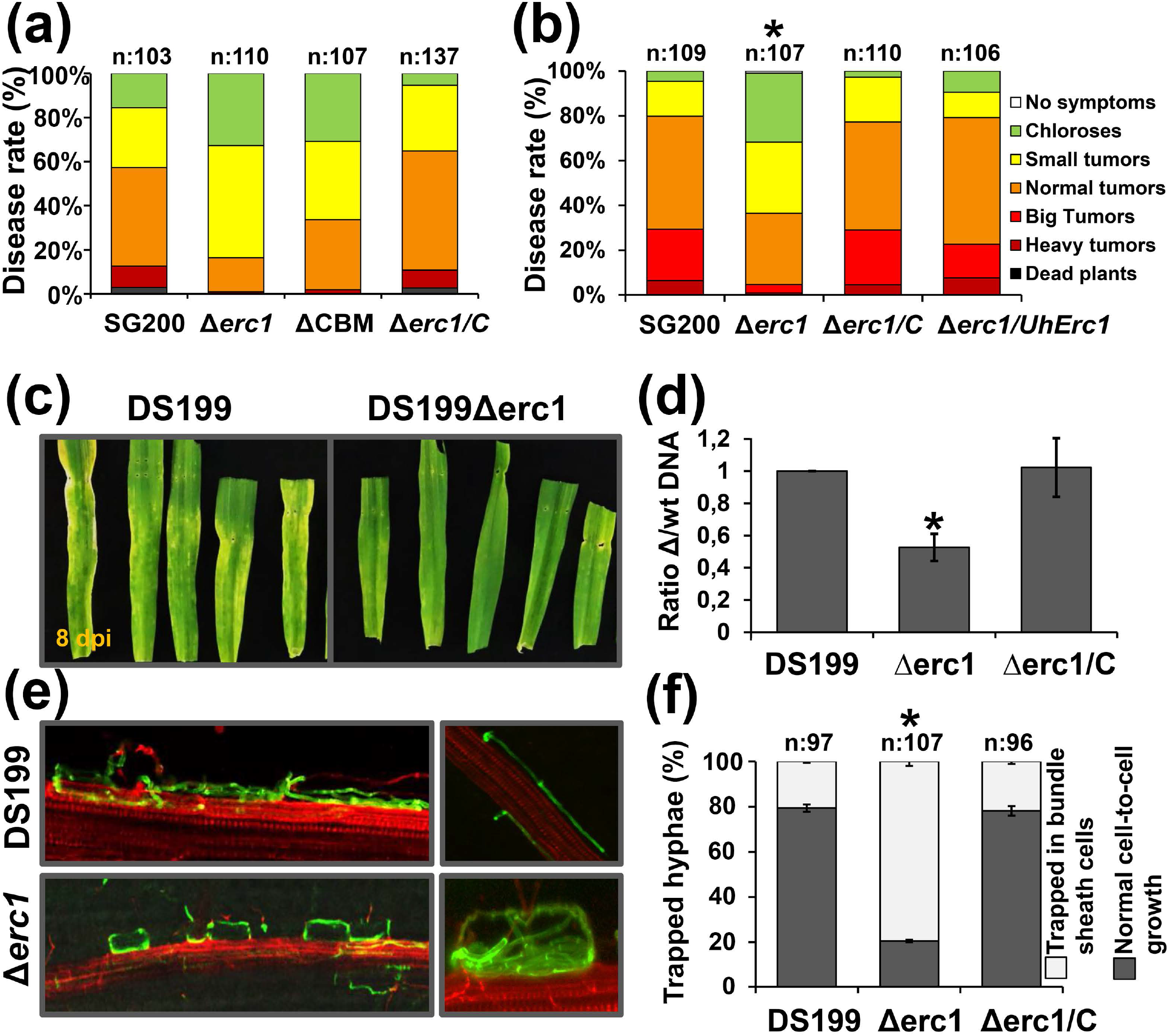
Erc1 is functionally conserved in smut fungi. **(a)** The carbohydrate binding module (CBM) is required for full function of *Ustilago maydis* Erc1. Disease assay was performed for *U. maydis* SG200, SG200Δerc1, SG200Δerc1/C strain, and SG200Δerc1/Erc1ΔCBM on EGB maize at 12 days post inoculation (dpi). Disease rates are given as a percentage of the total number of infected plants. n: number of infected maize seedlings. **(b)** Complementation of SG200Δerc1 mutant with *Ustilago hordei* UhErc1. Disease assay was performed for *U. maydis* SG200, SG200Δerc1, SG200Δerc1/C, and SG200Δerc1/UhErc1 strains on EGB maize plants at 12 days post inoculation (dpi). Three biological replicates were performed for all disease assays. n: number of infected maize seedlings. Asterisks above bars indicate significant differences (One-way ANOVA followed by Tukey multiple comparison test was performed, p<0.05). **(c)** UhErc1 is a virulence factor during barley infection. Disease assay was performed with *U. hordei* DS199 and DS199Δerc1 mutant strains on 13-day-old barley seedlings. Pictures were taken at 8 dpi. **(d)** Quantification of fungal biomass of DS199, DS199Δerc1 mutant and DS199Δerc1/C complementation strains on barley at 8 dpi. qPCR was performed to determine fungal biomass by using gDNA isolated from *U. hordei* DS199, DS199Δerc1 mutant and DS199Δerc1/C infected barley leaves. Three biological replicates were performed for all disease assays. n: number of infected barley seedlings. Asterisks above bars indicate significant differences (p<0.05, student t-test). **(e)** Microscopic observation of trapped *U. hordei* DS199 and DS199Δerc1 mutant hyphae in barley bundle sheath cells at 8 dpi via WGA-AF488/Propidium iodide staining. WGA-AF488 (green color - fungal cell wall): excitation at 488 nm and detection at 500-540 nm. PI (red color - plant cell wall): excitation at 561 nm and detection at 580–630 nm. **(f)** Quantification of cell-to-cell penetration efficiency of *U. hordei* DS199, DS1990Δuhafu1 mutant and DS199Δerc1/C strains. The graph depicts the total number of trapped *U. hordei* hyphae in barley bundle sheaths cells at 8 dpi. Three biological replicates were performed for the experiment. n: number of penetrated maize cells. Asterisks above bars indicate significant differences (p<0.05, student t-test).

### Erc1 is functionally conserved in covered smut of barley

Homology search and phylogenetic tree analysis show the presence of Erc1 in all available genomes of smut fungi **(Fig. S1 and Fig. S3)**. Considering the high degree of sequence conservation, we asked whether this effector could be functionally conserved in smut fungi, despite its highly specific virulence function in *U. maydis*. Thus, we expressed the *Erc1* gene from *Ustilago hordei*, the covered smut of barley, in the SG200Δerc1 mutant under the control of the native *U. maydis Erc1* promoter. Disease assays performed with the *U. maydis* SG200, SG200Δerc1 mutant, SG200Δerc1/Erc1 and SG200Δerc1/UhErc1 strains showed that UhErc1 fully restored virulence of the SG200Δerc1 mutant **(Fig. 2b)**. Complementary, we deleted the *UhErc1* gene in *U. hordei* to test its contribution to fungal virulence during barley colonization. The inoculation of barley seedlings with *U. hordei* strains DS199, DS199Δerc1 and DS199Δerc1/Erc1 showed significantly reduced fungal biomass of DS199Δerc1 compared to DS199 and the DS199Δerc1/Erc1 complementation strains **(Fig. 2c-d)**. Moreover, the DS199Δerc1 mutant showed a cell-arrest phenotype in the bundle sheath cells in barley leaves, mirroring the situation in *U. maydis* **(Fig. 2e-f)**. 80% of cell-to-cell movement attempts of the DS199Δerc1 mutant failed in bundle sheath cells compared to 20% for both DS199 and DS199Δerc1/Erc1 complementation strains **(Fig. 2e-f)**. Together, these results demonstrate that the cell-type specific virulence function of *Erc1* is conserved among the maize pathogen *U. maydis* and the barley pathogen *U. hordei*.

### Erc1 is secreted to the biotrophic interface

To localize Erc1 during host colonization, Erc1 with a fused C-terminal mCherry tag was expressed in the SG200Δerc1 mutant under the control of its native promoter. Confocal microscopy was performed using maize leaves inoculated with the Erc1-mCherry. Pit2-mCherry, which was previously shown to localize in the maize apoplast (Mueller *et al*., 2013), was used as a positive control for secretion and Int.mCherry (cytosolic mCherry) served as a negative control (Fig. 3a). Confocal microscopy confirmed that both Erc1-mCherry and Pit2-mCherry showed fluorescent signals on the outer surface of fungal hyphal tips, while the Int.mCherry showed fluorescent signals only inside the fungal cell (Fig. 3a). After plasmolysis with 1 M sodium chloride solution, the Erc1-mCherry signals accumulated in the apoplastic space and at the site of cell-to-cell passage on the plant cell wall, indicating extracellular secretion of Erc1-mCherry (Fig. 3a). For a more detailed subcellular localization of Erc1, we performed transmission electron microscopy (TEM) of immunogold labelled maize leaf sections infected by an Erc1-HA expressing SG200 strain (Fig. 3b). A SG200 strain expressing secreted GFP-HA was used as a control to compare signal distribution and specificity. TEM micrographs obtained from immunogold labelling with a monoclonal antibody for HA depicted secretion of Erc1-HA to the biotrophic interface. Erc1-HA and GFP-HA signals were detected both in the fungal cell wall and in adjacent plant cell wall regions (Fig. 3b). There was no significant difference observed in signal distribution of Erc1-HA and GFP-HA observed, which confirms secretion and apoplastic localization of Erc1 and does not indicate any specific focal accumulation of the effector apart from its apoplastic localization.

**Fig. 3.**
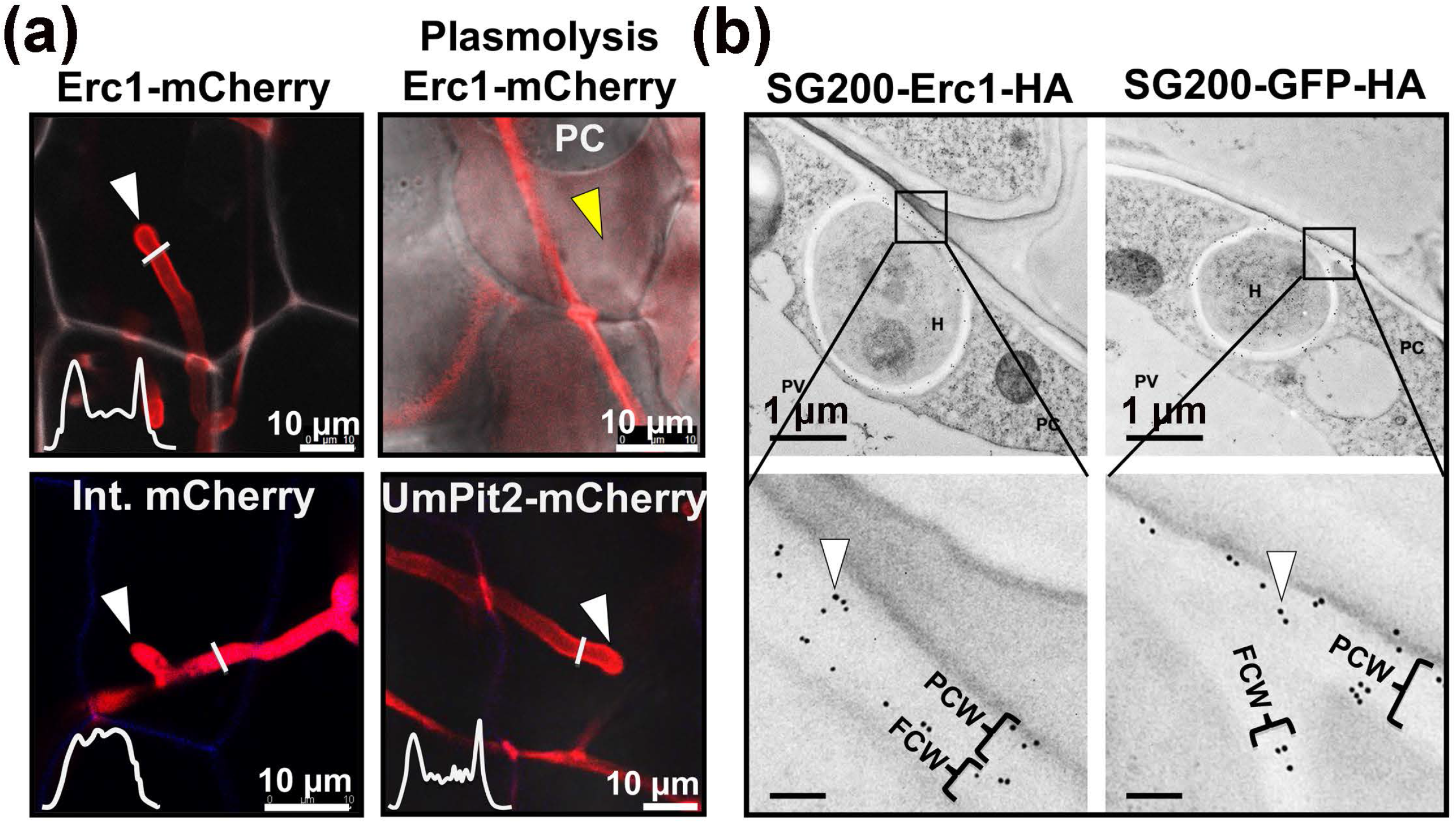
Localization of Erc1 in *Ustilago maydis* SG200 during maize colonization. **(a)** Erc1-mCherry was heterologously expressed in *U. maydis* SG200 strain under control of the native promotor with predicted native signal peptide for extracellular secretion. The SG200 strains expressing the Erc1-mCherry, UmPit2-mCherry (as a positive control for secretion) and cytosolic mCherry (as a negative control for secretion) were inoculated on maize seedlings and at 4 dpi confocal microscopy was performed to monitor the localization of each recombinant proteins. While both Erc1-mCherry and UmPit2-mCherry are secreted around the tip of the invasive hyphae, internal mCherry localizes to the fungal cytoplasm. The white graphs indicate the mCherry signal intensity along the diameter of the hyphae (illustrated by white lines in the image). White arrowheads indicate fungal hyphal tips and yellow arrowhead indicates apoplastic fluid after plasmolysis. Plasmolysis was performed with 1 M NaCl solution. **(b)** Transmission electron micrographs after immunogold labeling of secreted Erc1-HA and GFP-HA with a monoclonal antibody recognizing HA epitopes. Black dots depicted with white arrowheads indicate secretion of Erc1-HA to the biotrophic interface. FCW: Fungal cell wall, H: Hyphae, P: Plant cell cytoplasm, PCW: Plant cell wall.

### Erc1 does not exhibit α-L-arabinofuranosidase activity

To biochemically characterize the *U. maydis* Erc1 protein, N-terminally His tagged and C-terminally Myc/His-double-tagged Erc1 and its predicted active sites mutant (Erc1^M2X^: 410-412^GNE>AAA^ and 499^E>A^) were produced in the *Pichia pastoris* protein expression system. Subsequently, the recombinant proteins were purified via Nickel-NTA affinity chromatography (Fig. 4a). On SDS-PAGE, both recombinant proteins show two bands with higher molecular weight than the expected suggesting post-translational modification of these proteins **(Fig. 4a)**. A western blot analysis of the recombinant His-Erc1-Myc-His protein with antibodies specific for the His-tag and Myc-tag showed that, while the upper protein band is detectable both with an α-His and α-Myc antibody, the lower protein band is detectable only with an α-His antibody indicating a C-terminal cleavage of the tags **(Fig. 4a)**.

**Fig. 4.**
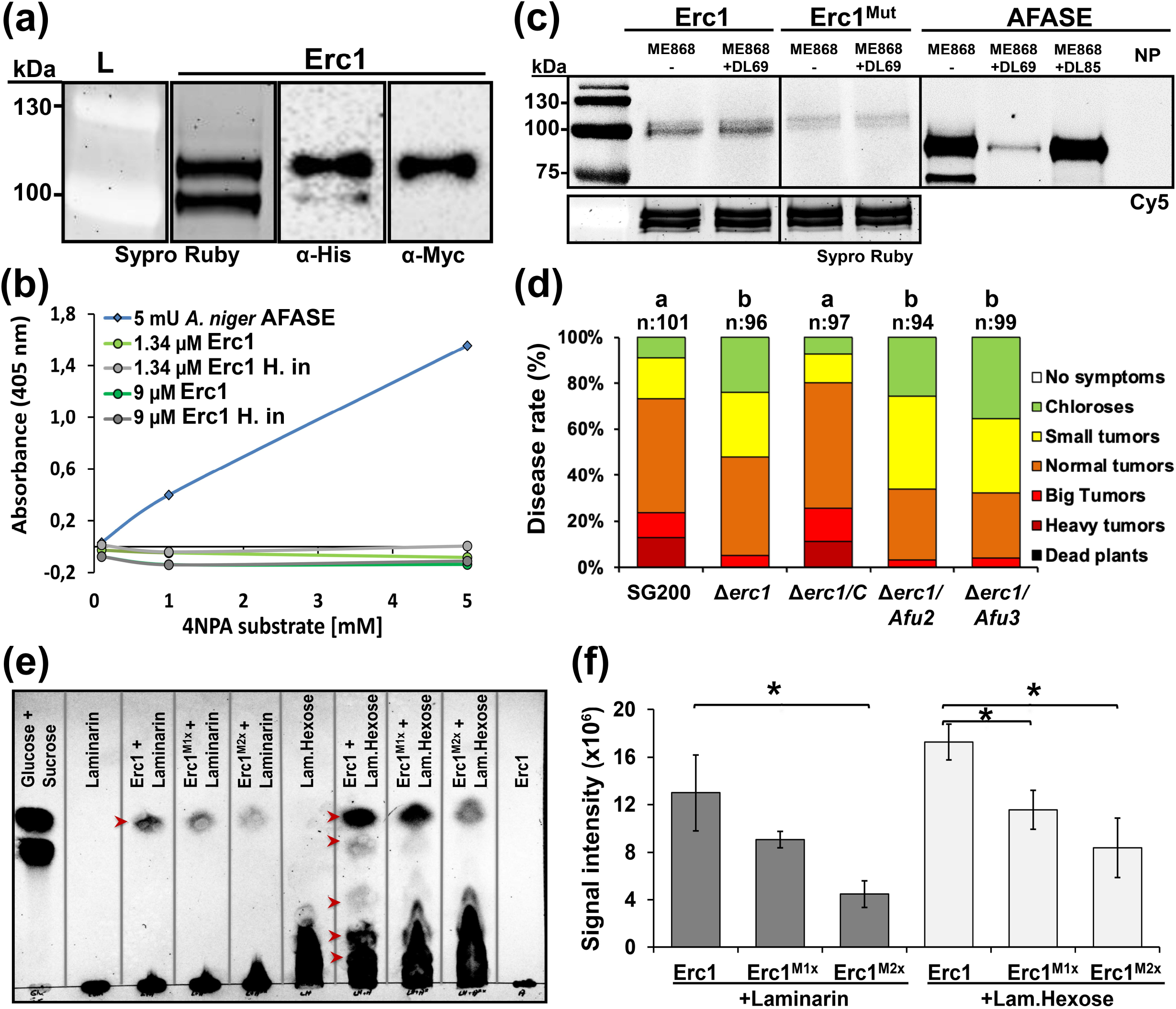
Functional characterization of the Erc1 protein. **(a)** Purification of His-Erc1-Myc-His recombinant protein. Sypro Ruby staining was performed to visualize Erc1 recombinant protein. Western blot (WB) analysis was performed with α-His- and α-Myc-specific antibody to detect His-Erc1-Myc-His protein in PVDF membrane. While two bands were detectable in Sypro Ruby staining and WB performed with α-His, only one band was detectable α-Myc-specific antibody. **(b)** α-L-arabinofuranosidase activity assay with 4-nitrophenyl α-L-arabinofuranoside (4NPA) substrate. A commercial α-L-arabinofuranosidase from *Aspergillus niger* (AFASE) was used as a positive control. **(c)** Activity-based protein profiling (ABPP) assay for Erc1. The Erc1, Erc1^M2x^ and AFASE recombinant proteins were incubated with DL69 α-L-arabinofuranosidase specific inhibitor. Plus (+) and minus (-) indicate the addition and the absence of the inhibitor, respectively. α-L-arabinofuranosidase specific ME868 probe was added as indicated (+/-). The probe was detected by scanning the in-gel fluorescence with Cy5 filter (Ex. 650 nm, Em. 670 nm). Protein of loaded samples were visualized via Sypro Ruby (Ex. 450 nm, Em. 610 nm). NP: no probe control. **(d)** Complementation of SG200Δerc1 mutant with *Afu2* and *Afu3* under the native *Erc1* promoter. Disease assay was performed for *Ustilago maydis* SG200, SG200Δerc1 mutant, SG200Δerc1/C, SG200Δerc1/Afu2 and SG200Δerc1/Afu3 strains on Early Golden Bantam (EGB) maize plants at 12 days post inoculation (dpi). Disease rates are given as a percentage of the total number of infected plants. Three biological replicates were performed for disease assay. n: number of infected maize seedlings. One-way ANOVA followed by Tukey multiple comparison test was performed, p<0.05. **(e-f)** Thin layer chromatography (TLC) assay was performed to demonstrate β-1,3-glucanase activity of Erc1 on laminarin and laminarihexaose substrate. Erc1, Erc1^M1x^ and Erc1^M2x^ recombinant protein were incubated with laminarin and laminarihexaose. Samples were loaded on TLC Silica gel 60 F_254_ plate. Glucose + Sucrose mix was used as reference. A mixture of n-propanol:ethanol:water (7:2:1/v:v:v) was used as a mobile phase. Laminarin, laminarihexaose and their hydrolysis products were visualized by spraying the TLC plate with detection solution. **(f)** The signal intensity of bands represent glucose was quantified by using ChemiDoc Bio-Rad imaging machine. Three biological replicates were performed and asterisks above bars indicate significant differences (p<0.05, student t-test).

Moreover, although both lower and upper Erc1 bands occur in similar protein amounts in a Sypro Ruby stained gel, the signal intensity of the upper band with the α-His antibody was much higher than the lower band, indicating also partial N-terminal cleavage **(Fig. 4a)**.

To test for the predicted α-L-arabinofuranosidase activity of Erc1, an enzyme activity assay was carried out with 4-nitrophenyl-α-L-arabinofuranose as a substrate. This substrate was incubated with both *U. maydis* Erc1 and a commercial arabinofuranosidase from *Aspergillus niger* (AFASE) as a positive control. While the commercial AFASE enzyme exhibited the expected α-L-arabinofuranosidase activity, no activity was detected in the samples incubated with the *U. maydis* Erc1 recombinant protein **(Fig. 4b)** suggesting that under these tested conditions and with this particular substrate Erc1 was not acting as an α-L-arabinofuranosidase. In addition, an activity-based protein profiling (ABPP) assay was performed by using the α-L-arabinofuranosidase specific fluorescent probe (ME868) and two specific inhibitors (DL69 and DL85) as controls **(Fig. 4c)** (McGregor *et al*., 2020). In the ABPP assay, the fluorophore-tagged probe binds covalently and specifically to the active site of the enzyme. The fluorescent signal levels indicate enzymatic activity, while lower/no fluorescent signal in the presence of both hydrolase-specific probe and inhibitor shows inhibition of a specific enzymatic activity. The ABPP assay showed that the ME868 probe can specifically bind to the AFASE enzyme control, and pre-treatment of AFASE with α-L-arabinofuranosidase-specific DL69 and DL85 inhibitors prevent this binding demonstrating the specificity of the assay and confirming the α-L-arabinofuranosidase activity of AFASE **(Fig. 4c)**. However, compared to AFASE only a very weak signal was observed for Erc1, which could not be inhibited by the α-L-arabinofuranosidase-specific inhibitors and therefore the weak signal can be considered as an unspecific background signal **(Fig. 4c)**. In addition to the enzyme activity assays, we complemented the SG200Δerc1 mutant with *Afu2* and *Afu3* (encoding the commercially available α-L-arabinofuranosidase) being expressed under the control of the native *Erc1* promoter. The resulting strains were used to test whether an enzyme with α-L-arabinofuranosidase activity could rescue the virulence phenotype of the *erc1* deletion mutant. However, disease assays performed with *U. maydis* strains SG200, Δerc1 mutant, Δerc1/C, Δerc1/Afu2 and Δerc1/Afu3 complementation strains showed that neither Afu2 nor Afu3 can complement the virulence defect of the Δerc1 mutant **(Fig. 4d)**.

### Erc1 exhibits exo-β-1,3-glucanase activity

Since α-L-arabinofuranosidase activity of Erc1 could not be demonstrated and none of the two other *U. maydis* α-L-arabinofuranosidases could restore the Δ*erc1* mutant infection phenotype, we decided to test whether Erc1 might exhibit another hydrolase activity such as endoglucanase activity being reported for some GH51 hydrolases. To this end, we incubated the recombinant Erc1 protein with several polysaccharides including β-1,4-glucan, laminarin, lichenan, xylan and arabinoxylan and used thin-layer chromatography (TLC) to visualize the released sugars from the tested candidate polymers. This approach revealed that Erc1 hydrolyses laminarin, which consists of a linear β-1,3-glucan with β-1,6-branches. Thus, Erc1 exhibits β-glucanase activity **(Fig. S4a)**. Laminarin polysaccharides consist of 20-25 units of linear β-1,3-glucan with β-1,6-linkages. To test whether Erc1 cleaves the β-1,3-glucan backbone or the β-1,6-linked branches, we performed an assay using laminarihexaose, which contains only linear β-1,3-glucan linkages **(Fig. 4e-f)**. The TLC results showed that Erc1 can hydrolase laminarihexaose demonstrating a β-1,3-glucanase activity. Furthermore, TLC performed with laminarin substrate incubated with Erc1, commercial exo-β-glucanase and endo-β-glucanase enzymes showed that, unlike the commercial endo-β-glucanase, which released several products with different intermediate sizes, both Erc1 and the exo-β-glucanase enzymes released only single glucose-moieties, suggesting an exo-β-1,3-glucanase activity of Erc1 **(Fig. S5a)**. To confirm the predicted active sites of Erc1, single (Erc1^M1x^: 410-412^GNE>AAA^) and double (Erc1^M2x^: 410-412^GNE>AAA^ and 499^E>A^) active site-mutated recombinant Erc1 proteins were incubated with both substrates. TLC showed a reduced enzymatic activity compared to recombinant wild-type Erc1. The residual enzymatic activity of the mutated Erc1 suggests the presence of additional active site/s, which we could not identify in this study **(Fig. 4e-f)**.

### Erc1 binds to plant cell wall components and suppresses laminarihexaose-induced ROS burst in plant leaves

To test the function of the predicted carbohydrate binding module (CBM) of Erc1, we performed carbohydrate binding assays. Recombinant Erc1 protein was incubated with insoluble polysaccharides originating from either the fungal cell wall (chitin and chitosan) or the plant cell wall (cellulose, xylan, β-1,4-glucan and lichenan). The subsequent pull-down assay showed that Erc1 did not bind to any of the tested fungal cell wall components **(Fig. S5b)**. In contrast, it bound to the plant cell wall components cellulose and lichenan, suggesting that Erc1 primarily binds the plant cell wall **(Fig. S5b)**. Co-incubation of Erc1 with either cellulose or lichenan did not produce any hydrolyzed products indicating that both polymers do not represent enzymatic substrates of Erc1 **(Fig. S4a)**. Monosaccharide analysis of the matrix polymers present in maize leaves treated with either mock, SG200, or SG200Δerc1 *U. maydis mutants* at 9 dpi did show differences between mock and SG200 infected leaf tissues as the wall arabinose content was increased, whereas the wall matrix glucose was decreased, but no differences in plant wall composition were observed between SG200 and SG200Δerc1 inoculations **(Fig. S4b)**. These data indicate that Erc1 does not quantitatively hydrolyze polymers in the plant cell wall.

Next, we performed a ROS-burst assay using cellotetraose as an elicitor to test whether Erc1 might interfere with cellotetraose-mediated accumulation of reactive oxygen species in barley leaf discs. Cellotetraose or chitosan (as a positive control) were pre-incubated with or without recombinant Erc1 and tested for their ability to induce a ROS burst on barley leaf discs **(Fig. S5c)**. The ROS production of chitosan alone showed the typical ROS burst with a peak occurring at 10 minutes after incubation **(Fig. S5c)**. Similar to chitosan, cellotetraose alone also triggered the production of ROS, although the peak of ROS production was detected after 20-30 minutes incubation **(Fig. S5c)**. Erc1 alone also induced a ROS burst on barley leaf discs, but the addition of either chitosan or cellotetraose samples to Erc1 protein resulted in two-fold and five-fold increase in ROS burst levels, respectively **(Fig. S5c)**.

It is well documented that β-1,3-glucans, such as laminarin and laminarihexaose, can be recognized by plant cells to induce defense responses including a ROS burst (Wanke *et al*., 2020; Chandrasekar *et al*., 2022). To test whether Erc1 interferes with laminarihexaose-induced plant defenses, we performed laminarihexaose-induced ROS burst assays on barley leaf discs **(Fig. 5a-b)**. Laminarihexaose alone induced a stable ROS burst in the barley leaf discs but this burst was significantly reduced by addition of Erc1 protein **(Fig. 5a-b)**. In contrast, laminarihexaose incubated with Erc1^M2x^ did not cause a significant difference in the laminarihexaose-induced ROS burst. This indicates that i) enzymatic activity of Erc1 is required for suppression of the laminarihexaose-induced ROS burst and ii) that the residual activity observed for Erc1^M2x^ in the recombinant enzyme assay was not sufficient to block the laminarihexaose-mediated ROS burst **(Fig. 5a-b)**. In addition, we observed that both Erc1 and Erc1^M2x^ recombinant proteins alone induced a weak ROS burst in barley leaf discs **(Fig. 5a-b)**. While Erc1^M2x^ appeared to have an additive effect on the laminarihexaose-induced ROS burst, the Erc1 plus laminarihexaose reaction mixture resulted in a residual ROS accumulation similar to the level of induction caused by Erc1 alone, which suggests that the glucan-induced burst was completely abolished **(Fig. 5a-b)**. Complementary to the ROS burst assay in leaf discs, we performed RT-qPCR analysis on *PR*-gene expression in maize leaves infected with *U. maydis* SG200 and SG200Δerc1 strains at 4 dpi **(Fig. 5c)**. The RT-qPCR data revealed a significant increase in the expression level of *PR1, PR3, PR5, PRm6b* and *PR10* genes in maize leaves infected with *U. maydis* SG200Δerc1 compared to SG200-infected leaves **(Fig. 5c)**, indicating that Erc1 is required for prevention/suppression of host immune responses during host colonization. These data suggest that a conserved secreted Erc1 effector is an enzyme required for hydrolyzing fungal wall-derived β-1,3-glucans, so that these DAMP molecules will not be perceived by the host to induce ROS burst and expression of PR genes.

**Fig. 5.**
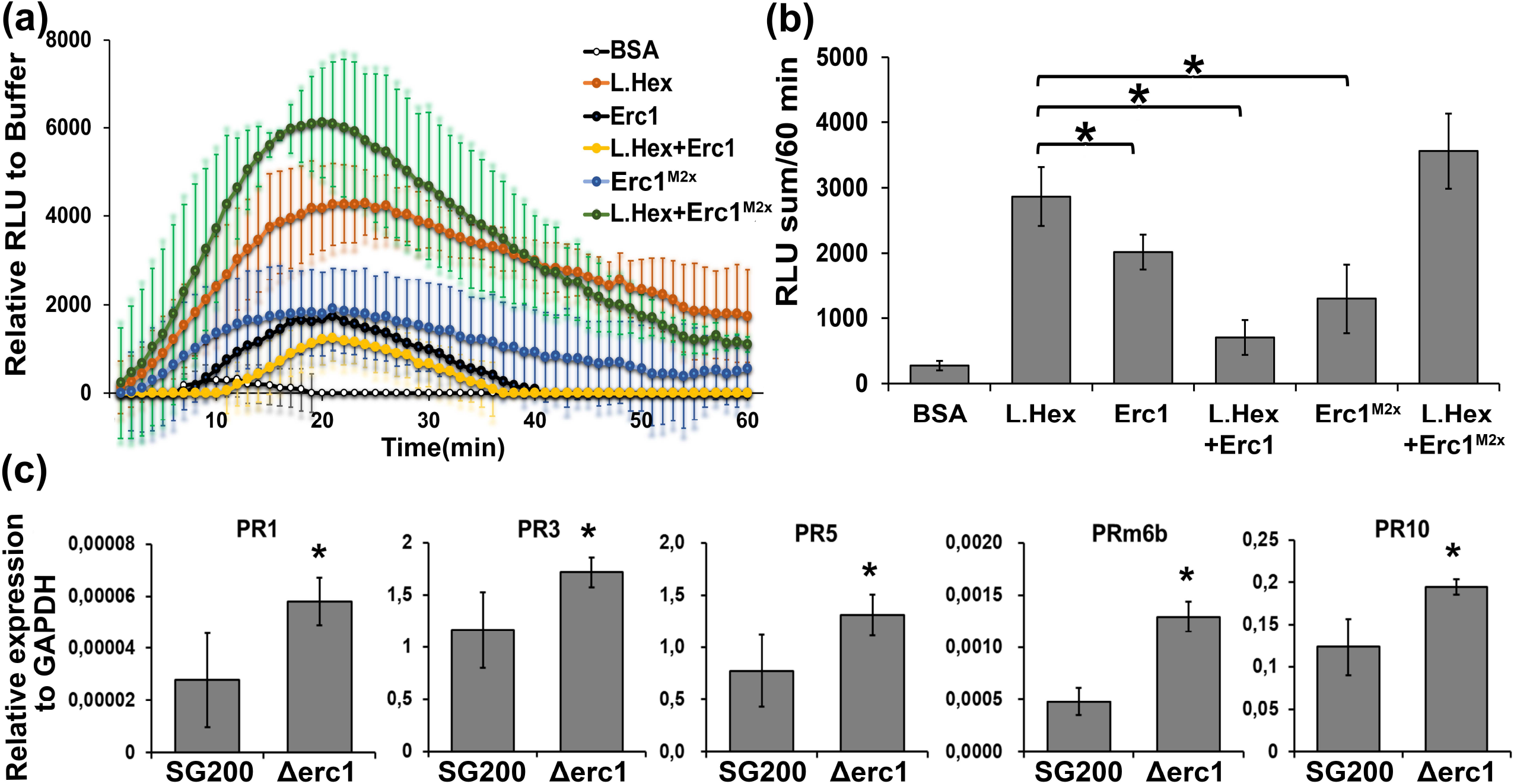
Smut Erc1 prevents induction of host defenses. **(a-b)** A ROS-burst assay was performed with barley leaf disks incubated with laminarihexaose), Erc1, Erc1^M2x^, and laminarihexaose incubated with Erc1 recombinant proteins. BSA was used as a negative control for background signal. Relative luminescense units (RLU) indicates ROS burst activity of treated barley leaf discs. The RLU are normalized with the buffer control. Each curves shows the mean of at least 9 technical leaf discs with three biological replicates. The error bars show the standard deviation of the three replicates. **(b)** Statistical analysis was performed with sum of RLU values of each sample. Asterisks above bars indicate significant differences (p<0.05, student t-test). **(c)** RT-qPCR for *pathogenesis related (PR)* gene expression on SG200 and SG200Δerc1 mutant strains infected maize leaves at 4 days post infection (dpi). The expression levels of maize *PR* genes, including *PR1, PR3, PR5, PRm6b* and *PR10*, were calculated relative to the *GAPDH* gene of maize. Asterisks above bars indicate significant differences (p<0.05, student t-test).

## Discussion

The corn smut fungus *U. maydis* infects all aerial organs of maize plants, including tassel, ears and leaves (Zuo *et al*., 2019). It has been shown that *U. maydis* deploys diverse sets of effectors to colonize different maize organs and around 45% of secreted proteins in *U. maydis* behave in an organ-specific manner (Skibbe *et al*., 2010; Schilling *et al*., 2014). However, there is little information about what determines the organ-specific function of *U. maydis* effectors. *U. maydis* Erc1 has previously been identified as an organ-specific virulence factor, which contributes to tumor formation in maize seedling leaves, but not in the tassel (Schilling *et al*., 2014). In this study, we provide a functional characterization of Erc1, a conserved core effector of smut fungi whose β-1,3 glucanase activity is required for its cell type-specific virulence function.

### *Ustilago maydis* α-L-arabinofuranosidases

Erc1 consists of a N-terminal signal peptide for extracellular secretion, a N-terminal carbohydrate binding module (CBM) and a GH51 domain. Both confocal microscopy and immunogold labelling TEM micrographs confirmed the secretion of Erc1 into the biotrophic interface, which is in line with the presence of a signal peptide. Although TEM micrographs did not show any specific Erc1 accumulation in fungal or plant cell walls, carbohydrate-binding assays revealed that the protein binds to the plant cell wall polysaccharides cellulose and mix-linkage glucan (i.e. lichenan). This does not only confirm the predicted CBM domain of Erc1, but also suggests that it may target maize plant cell wall components rather than the fungal ones. In addition, the importance of the CBM domain for the virulence function of Erc1 further supports the hypothesis that plant cell wall components are the biologically relevant virulence targets of Erc1.

Homology search showed that *U. maydis* has two Erc1 homologs, including Afu2 (GH51 family) and Afu3 (GH62 family). All three genes are predicted to encode α-L-arabinofuranosidases and are expressed during host colonization, but not in axenic culture (Lanver *et al*., 2018). Very high and continuous expression patterns of Erc1 in both *U. maydis*-maize and *U. hordei-*barley pathosystems indicate that Erc1 is required during the whole colonization process of these smut fungi (Lanver *et al*., 2018; Ökmen *et al*., 2018). Indeed, deletion of *erc1* in *U. maydis* and *U. hordei* resulted in a significant reduction in fungal virulence, demonstrating that Erc1 is a virulence factor of both smut fungi. Unlike Erc1, neither Afu2 nor Afu3 are required for *U. maydis* virulence (Lanver *et al*., 2014). However, the previous report by Lanver *et al*. (2014) also reported that a single deletion of the *Erc1* (named as “*afg1*”) gene had no effect on virulence in maize, while deletion of all three predicted *U. maydis* α-L-arabinofuranosidases decreased virulence. We can only speculate why the virulence function of Erc1 had been overlooked by Lanver and colleagues (e.g. different maize growth conditions, inoculation method or disease scoring). Similar to our findings, also a recent report by Marin-Menguiano and colleagues reported a significant contribution of Erc1 in maize smut virulence (Marín-Menguiano *et al*., 2019).

### Erc1 is a core effector in Ustilaginomycotina

In a previous comparative genome and secretome analysis performed by Schuster *et al*. (2018), Erc1 (Afu1) has been classified as a core effector being highly conserved in smut fungi (Lanver *et al*., 2018; Schuster *et al*., 2018). Our phylogenetic tree analysis shows that, although both Afu2 and Afu3 are present in a wide range of bacterial and fungal species, Erc1 is exclusively present in the Ustilaginomycotina, i.e. the smut fungi. The biotrophic rust fungi, which also belong to the Basidiomycetes and have largely overlapping plant hosts with the smuts, have homologs of Afu2, but they do not possess Erc1 homologs.

While a commercially available *U. maydis* Afu3 can hydrolyze wheat arbino-xylan, suggesting an actual α-L-arabinofuranosidase activity of this enzyme, we could not confirm the predicted α-L-arabinofuranosidase function of Erc1, neither with α-L-arabinofuranosidase specific ABPP probes nor with α-L-arabinofuranosidase specific 4NP-arabinofuranoside substrate. Moreover, complementation of the SG200Δerc1 mutant with *Afu2* and *Afu3* under the control of the *Erc1* promoter could not rescue the decreased virulence phenotype of *erc1* deletion mutant, indicating different enzymatic activity/virulence functions of these three proteins. Taken together the phylogenetic restriction to the Ustilaginomycotina and the distinct enzymatic activity, Erc1 appears to be a virulence factor that specifically evolved in smut-host interactions where it functions as a conserved core effector.

### Erc1 effector is a cell-type specific effector and is required for fungal cell-to-cell movement in host bundle sheath cells

Smut fungi penetrate and colonize host vascular bundles by moving from cell-to-cell (Kämper *et al*., 2006; Lanver *et al*., 2018). Our microscopy showed that Δerc1 mutants fail in cell-to-cell movement specifically in host bundle sheath cells, demonstrating a cell-type specific function of Erc1. Previous studies reported developmental and structural differences, including cell wall thickness and composition, between different plant cell types (Dengler *et al*., 1996). Both WGA-AF488/PI staining and protoplasting assays, which are regularly performed with maize leaves in our laboratory, demonstrated that there are some differences in the cell wall composition of bundle sheath and mesophyll cells **(Fig. 6a-b)**. While in the WGA-AF488/PI staining assay only bundle sheath cells could be stained with propidium iodide (PI) **(Fig. 6A, left picture, red signal)**, the cell wall-degrading enzyme mixture used in the protoplasting assay has an effect only on mesophyll cells, while bundle sheath cells remain intact **(Fig. 6b)**. We hypothesize that differences in cell wall composition or thickness might be the cause for the cell-type specific function of Erc1. In support of this, a WGA-AF488/PI staining assay performed with tassel tissue infected with *U. maydis* showed no PI staining, reflecting the absence of bundle sheath cells and similar cell wall composition with mesophyll cells **(Fig. 6a, red signal)**. In this regard, we suggest that the organ-specific function of Erc1 simply results from the absence of bundle sheath cells in maize tassel tissue.

**Fig. 6.**
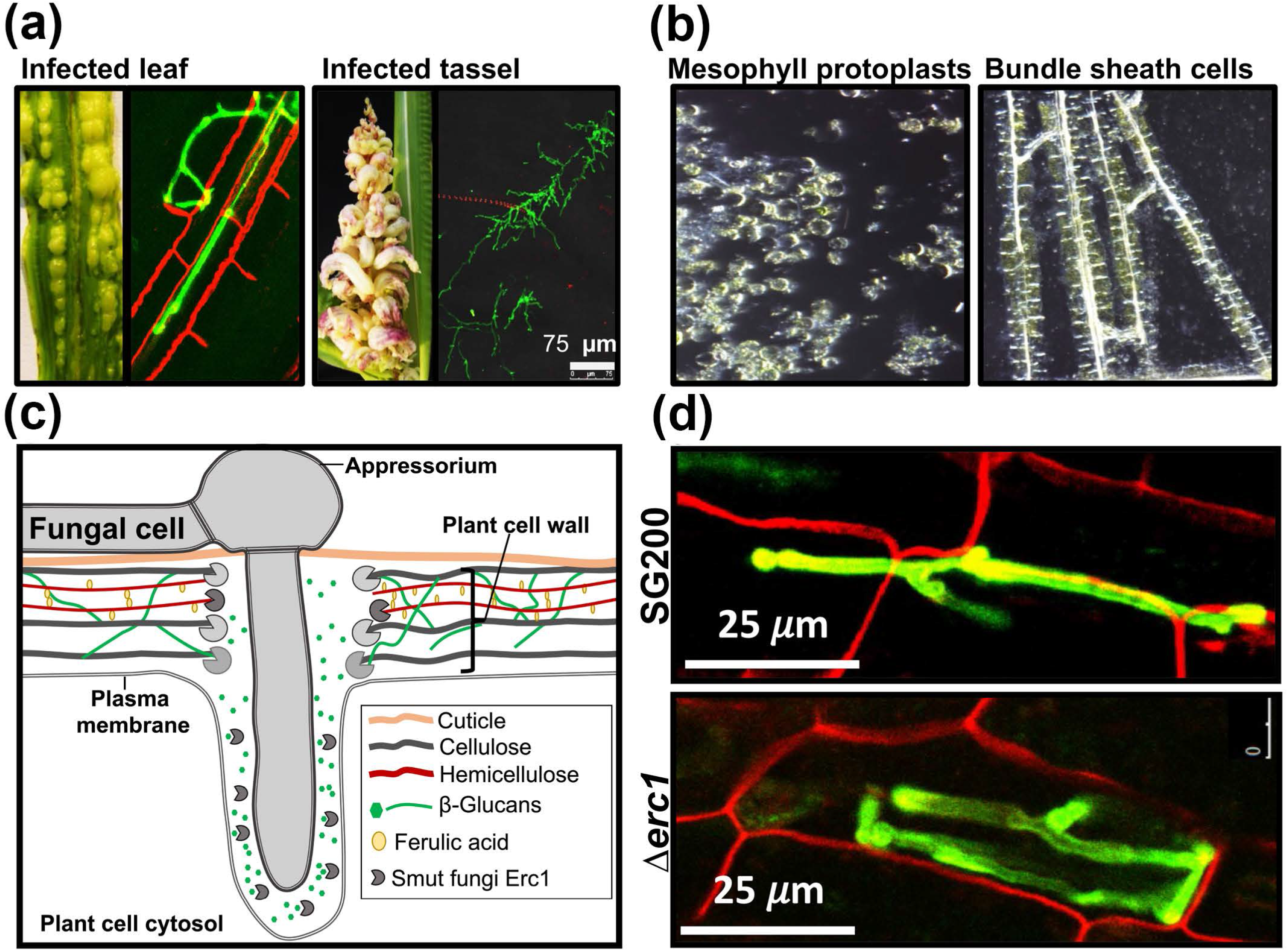
Cell type-specific function of Erc1. **(a)** Microscopic observation of trapped *Ustilago maydis* infected maize leaf and tassel tissues via WGA-AF488/Propidium iodide staining. WGA-AF488 (green color indicates fungal cell wall): excitation at 488 nm and detection at 500-540 nm. PI (red color indicates bundle sheath cells): excitation at 561 nm and detection at 580–630 nm. Unlike leaf tissue, no bundle sheaths cells are detectable in tassel tissue. **(b)** Protoplastation of maize leaf cells. Pictures were taken after the protoplasting process after treatment with plant cell wall degrading enzymes. While the used protoplastation enzyme mix has the ability to convert mesophyll cells into protoplast, bundle sheath cells remain intact. **(c)** Schematic depiction of the biotrophic interface of host-smut interaction. Fungus-derived proteins are depicted in gray. During host colonization, 1,3-β-glucan molecules are accumulated at the biotrophic interface and Erc1 may hydrolyze these 1,3-β-glucans to prevent the accumulation and subsequent recognition of DAMP molecules by the host plant. **(d)** While wild-type smut fungi have the ability to move from cell-to-cell in bundle sheath cells, Δerc1 mutants do not have the ability to fully suppress host immunity leading to the described cell-arrest phenotype (Modified from **Fig. 1c**).

### The enzymatic function of Erc1

It has been shown in many cases that fungal-derived extracellular GH family members and glucan-binding proteins play important roles in host penetration. While necrotrophs have relatively higher number and variation in their PCWDE repertoires to directly macerate and kill their host cells for nutrient acquisition, biotrophs fine-tune their limited number of PCWDEs to detoxify antimicrobial compounds, to acquire nutrients or to sequester MAMP molecules to prevent induction of host immunity (Bradley *et al*., 2022). For example, some of GH3 and GH10 family members are involved in detoxification of host-specific avenacin and α-tomatine, antimicrobial secondary metabolites from oat and tomato, respectively (Bowyer *et al*., 1995; Osbourn, 1996; Ökmen *et al*., 2013). Some GHs have the ability to sequester MAMP molecules released from fungal cell wall during host infection. A GH18 chitinase from *Magnaporthe oryzae* (MoChia1), as well as LysM lectins from a wide range of fungal pathogens bind and sequester chitin fragments released from the fungal cell wall, thereby preventing chitin-triggered immunity (de Jonge *et al*., 2010; Marshall *et al*., 2011; Mentlak *et al*., 2012; Yang *et al*., 2019).

We found that Erc1 hydrolyzes laminarin, which consists of 20-25 units 1,3-β-glucan with 1,6-β-glucan linkages, as well as laminarihexaose, which consists of only 1,3-β-glucans. Release of only glucose monomers from laminarin, but not any other intermediate-size sugar molecules, suggests an exo-β-glucanase activity of Erc1. Site-directed mutagenesis of two predicted active sites of Erc1 in *U. maydis* resulted in a significant reduction in virulence compared to the SG200, supporting the relevance of these two active sites in the virulence function of Erc1. Furthermore, TLC with Erc1^M1x^ and Erc1^M2x^ recombinant proteins showed that the enzymatic activity of both single- and double-active site mutant was significantly lower compared to the wild-type protein. However, the remaining residual enzymatic activity for both mutant proteins suggests a yet undiscovered active site of Erc1.

1,3-β-glucans are well described elicitors, inducing basal defense responses such as callose (another 1,3-β-glucan) deposition at infection sites (Wanke *et al*., 2020; Wanke *et al*., 2021; Chandrasekar *et al*., 2022). Recently, we performed immune gold labelling of 1,3-β-glucans on *U. hordei* infected barley leaf sections; subsequent TEM analysis revealed a strong accumulation of 1,3-β-glucans at the biotrophic interface (Ökmen *et al*., 2021). Thus, one could hypothesize that Erc1 hydrolyzes 1,3-β-glucans to prevent the accumulation and subsequent recognition of these DAMP molecules at the biotrophic interface **(Fig. 6c)**. This would also be consistent with our observation that Erc1 significantly reduces the laminarihexaose-induced ROS burst and that this activity depends on its enzymatic activity. Moreover, the presence or absence of Erc1 did not change monosaccharide contents of maize leaf cell walls during *U. maydis* infection, which indicates that the main target of Erc1 is not the plant cell wall itself. Thus, we suggest that Erc1 hydrolyzes 1,3-β-glucans accumulated at the biotrophic interface and in the absence of Erc1 the accumulated 1,3-β-glucans induces defense responses that result in cell arrest phenotype observed for Δerc1 mutants **(Fig. 6d)**.

Although fungal-derived GH family members take important roles in fungal virulence, in some cases these proteins, or their released products, can be recognized as MAMPs. For example, a GH45 cellulase (EG1) from *Rhizoctonia solani, a* GH12 xyloglucanase (BcXyg1) of *Botrytis cinerea* and a GH11 xylanase (EIX) of *Trichoderma viride* are recognized as MAMPs, triggering plant cell death and other immune responses (Ron & Avni, 2004; Ma *et al*., 2015; Zhu *et al*., 2017; Guo *et al*., 2022). Oligosaccharides released by *Cladosporium fulvum* CfGH17-1 (1,3-β-glucanase) from tomato cell walls trigger cell death upon recognition as DAMPs (Ökmen *et al*., 2019). Besides the suppression of laminarihexaose-induced ROS burst, Erc1 might be recognized by the plant independent of its enzymatic activity. Thus, Erc1 has an additive effect on the ROS-burst inducing activity of chitosan and cellotetraose elicitors. This additive effect on the induction of ROS-burst implies that Erc1 and carbohydrate elicitors are recognized by different pathways. On the other hand, it might suggest that other effectors are involved in the suppression of Erc1-mediated defense responses. Thus, several questions remain to be answered in upcoming studies: What determines cell-type specificity of Erc1? What is the receptor that recognizes Erc1 to induce ROS burst, but also what is the evolutionary origin of Erc1 proteins and how are they involved in the pathogenic lifestyle of smut fungi, i.e. their ability to colonize vegetative plant organs?

In summary, we functionally characterized a cell-type specific Erc1 effector from smut fungi that exhibits 1,3-β-glucanase activity and thus it is also involved in suppression of 1,3-β-glucan-mediated ROS burst in the host plant. Erc1 is functionally conserved in all plant pathogenic smuts and is required for cell-to-cell movement in bundle sheath cells.

## Materials and Methods

### Growth conditions for fungal and bacterial cultures

Growth conditions for fungal and bacterial strains used in this study are described in detail in **Supporting Information** part.

### Nucleic acids methods

Plasmid DNA from bacterial cells was isolated using the QIAprep Spin Miniprep Kit (Qiagen; Hilden, Germany) according to the manufacturer’s information. The genomic DNA isolation from *U. maydis* and *U. hordei* was performed according to the protocol described by Schultz *et al*. (1990). The polymerase chain reaction (PCR) utilizing Phusion© polymerase enzyme (Thermo Fisher Scientific;Darmstadt, Germany) was used to amplify specific DNA fragments by using gene specific primer pairs depicted in **Table S1**.

Total RNA isolation, cDNA synthesis and RT-qPCR analysis were described in **Supporting Information** part. The primers used for RT-qPCR are listed in **Table S1**.

### Construction of plasmids

All plasmid constructions were performed by using standard molecular biology methods according to molecular cloning laboratory manual of Sambrook *et al*. (1989). gDNA from *U. maydis* and *U. hordei* were used for the amplification of deletion and complementation constructs via PCR, whereas cDNA was used to amplify expression constructs. After PCR amplification of the desired genomic region with appropriate primers **(Table S1)**, the PCR fragments were digested with appropriate restriction enzymes and ligation reaction was performed with the T4-DNA ligase (New England Biolabs; Frankfurt a.M., Germany). All vector constructs, primer pairs and restriction sites that were used for the cloning procedures are indicated in **Table S1**. The nucleotide sequences of all constructs were confirmed via sequencing at the Eurofins sequencing facility (Germany).

Chemically competent *Escherichia coli* DH5α cells were transformed via heat shock assay according to standard molecular biology methods (Sambrook *et al*., 1989). *P. pastoris* was transformed as described in the yeast protocol handbook (Clontech, Mountain View, USA). Gene replacement and transformation assays for *U. maydis* and *U. hordei* protoplasts were conducted as described in Kämper (2004). All generated deletion and complementation mutant strains were confirmed via southern blot analysis for single integration event in the desired loci **(Fig. S6)**.

### CRISPR/Cas9 gene editing system

The CRISPR/Cas9-HF (high fidelity) gene editing system was used to knockout the *Erc1* gene in the *U. hordei* as described in Ökmen *et al*. (2021). To express sgRNA for the targeted gene, the *Ustilago maydis pU6* promotor was replaced with the *U. hordei pU6* promotor. The sgRNAs for knocking out the *U. hordei Erc1* gene was designed by E-CRISPR (http://www.e-crisp.org/ECRISP/aboutpage.html) **(Table S1)** (Heigwer *et al*., 2014). Plasmid construction for CRISPR/Cas9 was performed as described by Zou *et al*. (2020). The CRISPR/Cas9-HF vector was linearized with the restriction enzyme Acc65I, and subsequently assembled with spacer oligo and scaffold RNA fragment with 3’ downstream 20 bp overlap to the plasmid by using Gibson Assembly (Gibson *et al*., 2009).

### *Ustilago maydis* virulence assay

The *U. maydis* virulence assays on the EGB maize cultivar were performed as described in Redkar and Doehlemann (2016). Experimental details of disease assay for *U. maydis* and *U. hordei* are described in **Supporting Information** part.

### Heterologous protein production in *Pichia pastoris*

The *Pichia pastoris* KM71H-OCH protein expression system was used to produce N-terminally His and C-terminally Myc-His tagged *U. maydis* Erc1, Erc1^M1x^, and Erc1^M2x^ recombinant proteins. All genes for each respective protein were cloned into the pGAPZαA vector (Invitrogen; Carlsbad, USA) under the control of a constitutive promotor with an α-factor signal peptide for secretion. Protein expression and purification of recombinant Erc1 was described in detail in **Supporting Information** part.

### Enzyme activity assay and activity-based protein profiling

To determine the enzymatic activity of Erc1, 4-nitrophenyl α-L-arabinofuranoside (4NPA) activity and activity-based protein profiling (ABPP) assays were carried out. The commercially available arabinofuranosidase (AFASE) from *Aspergillus niger* (Megazyme, Ireland) was used as a positive control in both assays. Experimental details of both assays are described in **Supporting Information** part.

### Thin-layer chromatography (TLC) assay

To detect any carbohydrate hydrolysing activity of Erc1 a thin-layer chromatography assay (TLC) was performed with different polysaccharides incubated with purified Erc1 recombinant protein as described in Ökmen *et al*. (2019). Experimental details of TLC assay is described in **Supporting Information** part.

### WGA-AF488/Propidium iodide staining of infected plant tissues

The WGA-AF488 (Molecular Probes, Karlsruhe, Germany) and propidium iodide (Sigma-Aldrich) cell staining was performed according to Ökmen *et al*. (2018). WGA-AF488 stains fungal cell walls (green), while the propidium iodide stains plant cell walls (red). Experimental details of WGA-AF488/Propidium iodide staining is described in **Supporting Information** part.

### Localization of Erc1-mCherry with confocal microscopy

To visualize the secretion of Erc1-mCherry protein in *U. maydis* during maize colonization, the *U. maydis* SG200 strains expressing Erc1-mCherry, Pit2-mCherry and cytosolic mCherry were inoculated on maize seedlings. Subsequently, infected maize leaves were observed for localization of Erc1-mCherry at 4 dpi with a Leica confocal microscopy SP8. For mCherry fluorescence of hyphae in maize tissue, an excitation at 561 nm and detection at 580–630 nm were used. Plasmolysis was performed with 1 M NaCl solution dropped on the leaf sample.

### ROS burst assay

For ROS burst assay, 12 leaf disks, which were dissected from the second leaf of 13 days old Golden Promise barley cultivar, were incubated with 200 μl water in a 96 well plate (Thermo Fisher Scientific-Nunclon 96 Flat Bottom White Polystyrene) overnight in the dark. The next day the water was replaced with a 150 μl reaction mixture containing a luminol solution (100 μM L-012 and 20 μg ml^-1^ HRP). The following reaction mixtures were used in the ROS burst assay for barley leaf disks: (1) Buffer 20 mM KPO_4_ (pH6) (2) BSA, (3) 250 μM laminarihexaose, (4) 3 μM Erc1, (5) 250 μM laminarihexaose incubated with 3 μM Erc1, (6) 3 μM Erc1^M2x^, (7) 250 μM laminarihexaose incubated with 3 μM Erc1^M2x^. The luminescence from each well was measured with the following settings in a Tecan Infinite 200 Pro plate reader: Kinetic cycle: 60, interval time: 1 min, integration time: 450 ms.

### Transmission electron microscopy and immunogold labelling

#### High pressure freezing, freeze substitution and resin embedding

For TEM observations, 2 mm leaf discs from maize leaves infected with *U. maydis* expressing Erc1-HA and GFP-HA under the control of the native Erc1 promoter and signal peptide, were excised from 1 cm below infected area using a biopsy punch and processed by means of high pressure freezing (HPF) following the procedure described in Micali *et al*. (2011). Experimental detail of high pressure freezing and embedding is described in **Supporting Information** part.

#### Sectioning, immunogold labelling and transmission electron microscopy

Ultrathin (70-90 nm) sections were collected on nickel slot grids as described by Moran and Rowley (1987). Experimental detail of immunogold labelling is described in **Supporting Information** part.

### Plant cell wall analysis

Plant cell walls were isolated from mock, SG200 and SG200Δerc1 mutant treated Early Golden Bantam (EGB) maize leaves at 9 dpi. Inoculated plant tissues were lyophilized and homogenized using a MM400 mixer mill (Retsch Technology). Preparation of a destarched alcohol insoluble residue (cell wall preparation) was performed as described in Foster *et al*. (2010). The monosaccharide composition of that residue was analyzed via high-performance anion-exchange chromatography coupled with pulsed electrochemical detection (HPAEC-PAD), as previously described (Ramírez *et al*., 2018) using a 940 Professional IC Vario ONE/ChS/PP/LPG instrument (Metrohm) equipped with CarboPac PA20 guard and analytical columns. The monosaccharides were quantified based on known concentrations of standards.

### Bioinformatics methods

The SignalP 5 software program was used to predict an N-terminal secretion signal peptide within a protein sequence (http://www.cbs.dtu.dk/services/SignalP/). For protein domain prediction the website Pfam (http://pfam.xfam.org/) was used. A minimum evolution tree was constructed by using an alignment of the full-length amino acid sequence of Erc1 homologs obtained from the NCBI database for different microorganisms. The minimum evolution tree was constructed by using the software Mega7 (http://www.megasoftware.net/mega.php) with a minimum evolution algorithm performing 1000 bootstraps.

## Acknowledgments

The authors would like to thank Katharina Lufen, Institute for Plant Cell Biology and Biotechnology, Heinrich Heine University, Düsseldorf for excellent technical assistance with respect to the monosaccharide compositional analysis. GD and MP acknowledge support from the Cluster of Excellence on Plant Sciences (CEPLAS) funded by the Deutsche Forschungsgemeinschaft (DFG, German Research Foundation) under Germany’s Excellence Strategy – EXC 2048/1 – Project ID: 390686111. The authors would also like to thank Ila Rouhara from Central Microscopy, Max-Planck-Institute for Plant Breeding Research for excellent technical assistance with respect to TEM. The authors thank Prof. Dr. Hermen Overkleeft from Leiden Institute of Chemistry, LIC/Chemical Biology (The Netherlands) for providing ABPP probes and inhibitors.

## Author Contribution

B.□. and G.D. conceived the project. B.Ö., E.J., L.S., N.F., Y.J.L., and R.W. carried out transformation, disease assays, protein production, ROS and TLC assays; U.N. carried out the immunogold labeling followed by TEM assay; M.P. carried out cell wall analyses; B.Ö. wrote the manuscript with input from all authors.

## Supporting Information

### Supporting Figures

**Fig. S1. Amino acid alignment of Erc1 from different smut fungi. (a)** ClustalOmega and GeneDoc software programs were used to construct this alignment. The blue box represents the predicted signal peptide using SignalP5, the red box shows the predicted carbohydrate-binding domain (CBM) and the green box shows glycoside hydrolase family 51 (GH51) domain. The GNE and E, which are depicted above the amino acid sequence in red, are predicted active sites for Erc1. Mp: *Melanopsichium pennsylvanicum*, Uh: *Ustilago hordei*, Ub: *Ustilago bromivora*, Ph: *Pseudozyma hubeiensis*, Um: *Ustilago maydis* Sr: *Sporisorium reilianum*, Ssc: *Sporisorium scitamineum*. **(b) Schematic presentation of Erc1 (Afu1), Afu2 and Afu3**. SP: Signal peptide; CBM: Carbohydrate-binding module, GH: Glycoside hydrolase family.

**Fig. S2. (a)** Expression pattern of *GH51* genes, including *Erc1* (*Afu1*), *Afu2* and *Afu3*, in *Ustilago maydis* during maize infection. The transcriptome data created by Lanver *et al*. (2018) were used to create this bar graph (Lanver *et al*., 2018). *UmPit2* (as a high expressed effector) and *UmSee1* (as a low expressed effector) effector genes were also added to the graph for reference. **(b)** Afu2 and Afu3 are not required for full virulence of *U. maydis* during maize infection. Disease symptoms caused by *Ustilago maydis* SG200, SG200Δerc1, SG200Δerc1/UhErc1, SG200Δafu2-3 and SG200Δ3xafu/Erc1 on Early Golden Bantam (EGB) maize cultivar at 12 days post inoculation (dpi). Disease rates are given as a percentage of the total number of infected plants. Two biological replicates were performed for disease assay. n: number of infected maize seedlings.

**Fig. S3**. Phylogenetic tree analyses of GH51 proteins. A minimum evolution tree was constructed by using an alignment of the full-length amino acid sequence of Erc1 homologs obtained from the NCBI database for different microorganisms. The minimum evolution tree was constructed by using the software Mega7 by using the minimum evolution algorithm performing 1000 bootstraps. PHSY_003700: *Pseudozyma hubeiensis* SY62; SRS1_12911: *Sporisorium reilianum* f. sp. *reilianum*; sr12911: *Sporisorium reilianum* SRZ2; SSCI78560.1: *Sporisorium scitamineum*; PANT_22c00212: *Moesziomyces antarcticus* T-34; PSEUBRA_SCAF2g02877: *Kalmanozyma brasiliensis* GHG001; UBRO_02718: *Ustilago bromivora*; UHOR_02718: *Ustilago hordei*; BN887_04651: *Melanopsichium pennsylvanicum*; BCV70DRAFT_193121: *Testicularia cyperi*; PFL1_01248: *Anthracocystis flocculosa* PF-1; PHSY_001491: *Pseudozyma hubeiensis* SY62; PaG_05269: *Moesziomyces aphidis* DSM 70725; Moror_11954: *Moniliophthora roreri* MCA 2997; PYCCODRAFT_1357629: *Trametes coccinea* BRFM310; BOTBODRAFT_205067: *Botryobasidium botryosum* FD-172 SS1; GYMLUDRAFT_33770: *Gymnopus luxurians* FD-317 M1; RSOLAG22IIIB_03069: *Rhizoctonia solani*; ARMGADRAFT_1053725: *Armillaria gallica*; SCHPADRAFT_905275: *Schizopora paradoxa*; AFLA_015670: *Aspergillus flavus*; AFGD_006065: *Aspergillus flavus*; AO090012000298: *Aspergillus oryzae*; P875_00021913: *Aspergillus parasiticus* SU-1; M408DRAFT_30410: *Serendipita vermifera* MAFF 305830; Vi05172_g6833: *Venturia inaequalis*; OH76DRAFT_1461559: *Polyporus brumalis*; PENSUB_1664: *Penicillium subrubescens*; ZT3D7_G5366: *Zymoseptoria tritici* ST99CH_3D7; V565_067700: *Rhizoctonia solani* 123E; PSEUBRA_SCAF1g00656: *Kalmanozyma brasiliensis* GHG001; PAN0_002c0906: *Moesziomyces antarcticus*; SRS1_12124: *Sporisorium reilianum* f. sp. *reilianum*; sr12124: *Sporisorium reilianum* SRZ2; PANT_7c00355: *Moesziomyces antarcticus* T-34; UHOR_01245: *Ustilago hordei*; SPSC_03938: *Sporisorium scitamineum*; UBRO_01245: *Ustilago bromivora*; SSCI26350.1: *Sporisorium scitamineum*; MELLADRAFT_107307: *Melampsora larici-populina* 98AG31; SSCI70960.1: *Sporisorium scitamineum*; sr15193: *Sporisorium reilianum* SRZ2; SRS1_15193: *Sporisorium reilianum* f. sp. *reilianum*; PANT_9d00343: *Moesziomyces antarcticus* T-34; PaG_02267: *Moesziomyces aphidis* DSM 70725; PSEUBRA_SCAF21g03514: *Kalmanozyma brasiliensis* GHG001; UBRO_06745: *Ustilago bromivora*; PHSY_003373: *Pseudozyma hubeiensis* SY62; UHOR_06745: *Ustilago hordei*; WP_086671302.1: *Amycolatopsis pretoriensis*; WP_100808766.1: *Streptomyces* sp. M56; WP_093701673.1: *Streptomyces* sp. MnatMP-M27; WP_079255736.1: *Streptomyces* autolyticus; WP_026469352.1: *Amycolatopsis balhimycina*; EPX63559.1: *Cystobacter fuscus* DSM 2262; WP_184195820.1: *Armatimonas rosea*; AARAC_001847: *Aspergillus arachidicola*; ABOM_009987: *Aspergillus bombycis*; AO090701000885: *Aspergillus oryzae* RIB40; BCIN_14g05510: *Botrytis cinerea* B05.10; P875_00033707: *Aspergillus parasiticus* SU-1; AFGD_012181: *Aspergillus flavus*; AN7908.2: *Aspergillus nidulans* FGSC A4; TGAM01_v202157: *Trichoderma gamsii*; CSAL01_09190: *Colletotrichum salicis*; BFJ70_g6661: *Fusarium oxysporum*; FCULG_00009021: *Fusarium culmorum*; DOTSEDRAFT_63078: *Dothistroma septosporum* NZE10; CB0940_03346: *Cercospora beticola*; IQ06DRAFT_98845: *Stagonospora* sp. SRC1lsM3a; CC77DRAFT_752970: *Alternaria alternata*

**Fig. S4. (a) Thin-layer chromatography (TLC) assay** to show enzymatic activity of Erc1 on different polysaccharides, including cellulose, lichenan, cellotetraose, β-glucan, laminarin, xylan and arabinoxylan. A mixture of n-propanol:ethanol:water (7:2:1/v:v:v) was used as a mobile phase. Carbohydrates and their hydrolysis products were visualized by spraying the TLC plate with detection solution and subsequent drying at 100°C for approximately 15 min. Red arrow head indicates hydrolysis product of laminarin. Glucose + sucrose mix was used as reference. **(b) Monosaccharide composition of maize cell walls**. Plant cell walls were isolated from mock, SG200 and SG200Δerc1-treated EGB maize leaves. The isolated plant cell walls were analyzed for their arabinose, galactose, glucose and xylose content. Asterisks above bars indicate significant differences (p<0.05, student t-test).

**Fig. S5. (a)** Thin-layer chromatography (TLC) assay was performed to demonstrate exo-β-1,3-glucanase activity of Erc1 on laminarin. Glucose + sucrose mix was used as reference. A mixture of n-propanol:ethanol:water (7:2:1/v:v:v) was used as a mobile phase. Laminarin and its hydrolysis products were visualized by spraying the TLC plate with detection solution and subsequent drying at 100°C for approximately 15 min. Red arrow heads indicate hydrolysis products of laminarin. Commercial exo- and endo-β-1,3-glucanases were used as controls. **(b)** Carbohydrate binding assay for recombinant Erc1 protein. Erc1 protein was incubated with insoluble chitin, chitosan, cellulose, xylan, β-glucan and lichenan. Subsequently, supernatant and pellet phases were analysed for presence of Erc1 protein via SDS-PAGE followed by coomassie blue staining. S: supernatant, P: pellet **(c)** To check whether Erc1 sequesters cellotetraose in order to prevent its recognition as a DAMP, a ROS-burst assay was performed with barley leaf disks incubated with cellotetraose, Erc1 and cellotetraose incubated with Erc1 recombinant protein. Chitosan was used as a negative control. Relative luminescence units (RLU) indicate ROS burst activity of treated barley leaf discs. The RLU are normalized with the buffer control. The error bars show the standard deviation of the three replicates.

**Fig. S6. Southern Blot analysis to confirm a single insertion event in different SG200**Δ**erc1 complementation strains**. gDNA of both SG200 and complementation strains were digested with *Bam*HI or *Sac*I restriction enzyme. Cbx gene was used as a probe to detect single insertion event. Expected band sizes for wild-type, single and multiple integration events are depicted below the figure.

### Supporting Table

**TableS1. Primer and construct list**.

