## Supplemental Material for "A conserved enzyme of smut fungi facilitates cell-to-cell movement in the plant bundle sheath"

**Corresponding authors:**

**Supporting Information: Figs S1-S6 and Table S1**

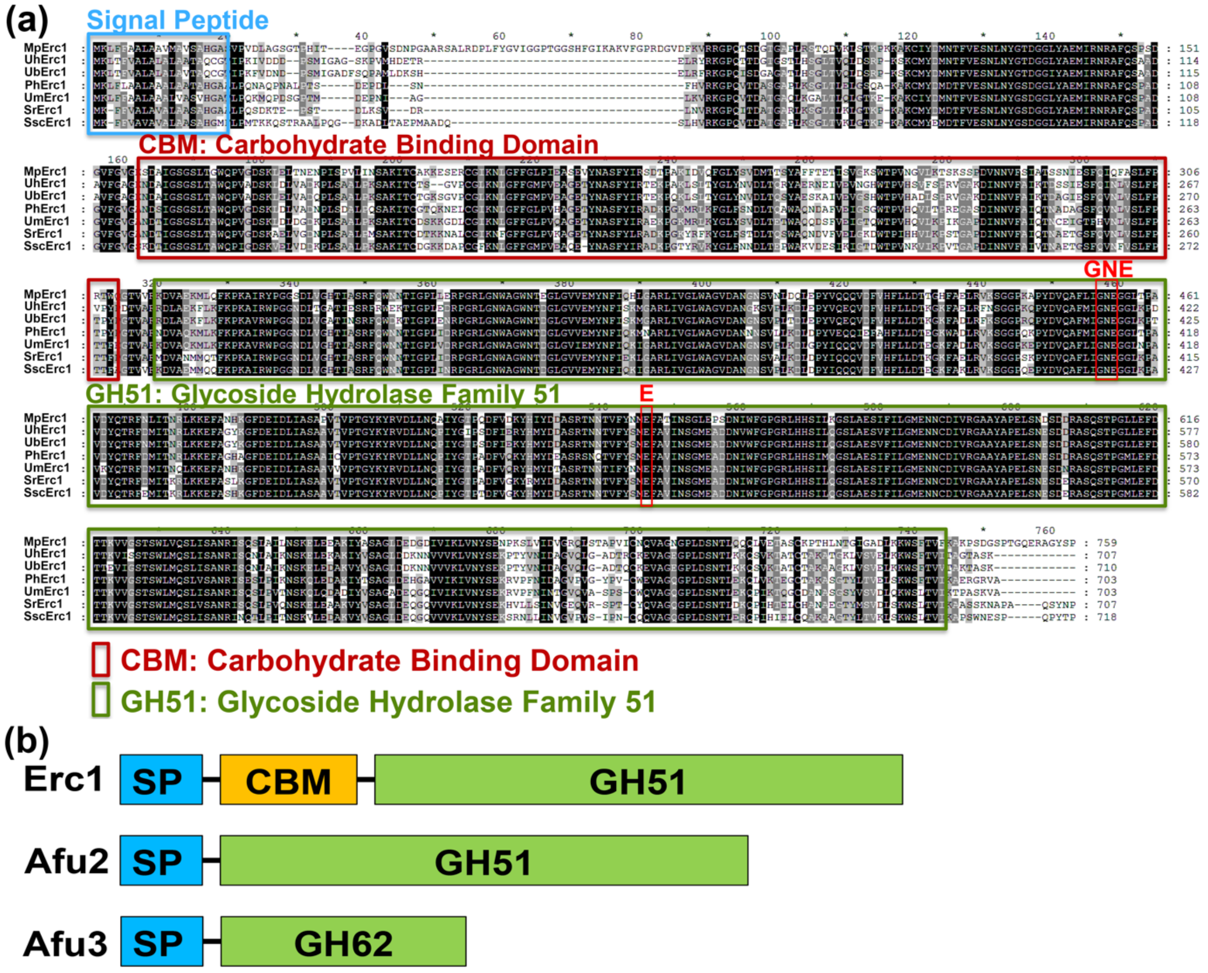


**Fig. S1. Amino acid alignment of Erc1 from different smut fungi. (a)** ClustalOmega and GeneDoc software programs were used to construct this alignment. The blue box represents the predicted signal peptide using SignalP5, the red box shows the predicted carbohydrate-binding domain (CBM) and the green box shows glycoside hydrolase family 51 (GH51) domain. The GNE and E, which are depicted above the amino acid sequence in red, are predicted active sites for Erc1. Mp: *Melanopsichium pennsylvanicum,* Uh: *Ustilago hordei*, Ub: *Ustilago bromivora*, Ph: *Pseudozyma hubeiensis*, Um: *Ustilago maydis* Sr: *Sporisorium reilianum,* Ssc: *Sporisorium scitamineum.* **(b) Schematic presentation of Erc1 (Afu1), Afu2 and Afu3.** SP: Signal peptide; CBM: Carbohydrate-binding module, GH: Glycoside hydrolase family.

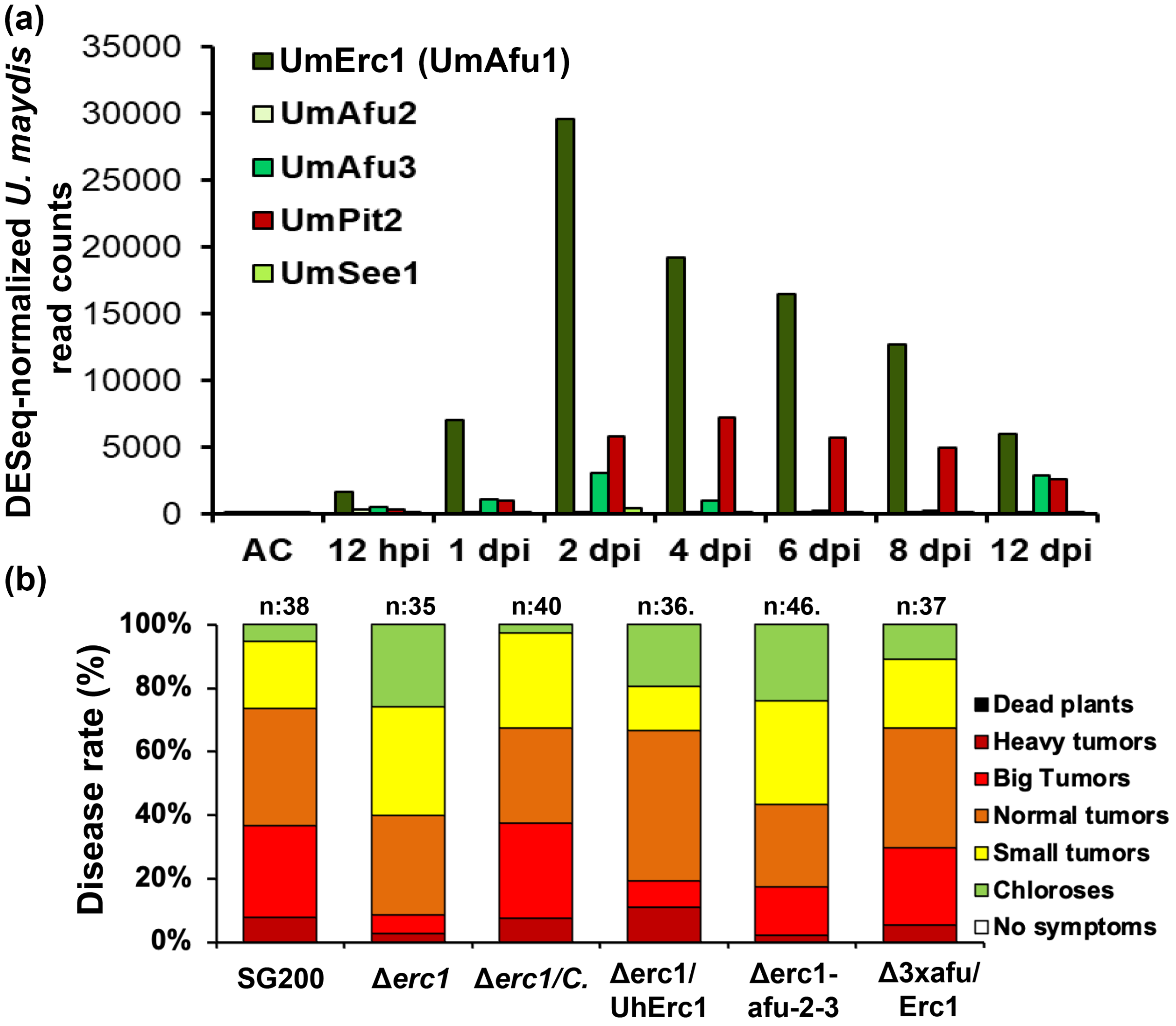


**Fig. S2.** **(a)** Expression pattern of *GH51* genes, including *Erc1* (*Afu1*), *Afu2* and *Afu3*, in *Ustilago maydis* during maize infection. The transcriptome data created by Lanver *et al*. (2018) were used to create this bar graph (Lanver *et al.*, 2018). *UmPit2* (as a high expressed effector) and *UmSee1* (as a low expressed effector) effector genes were also added to the graph for reference. **(b)** Afu2 and Afu3 are not required for full virulence of *U. maydis* during maize infection. Disease symptoms caused by *Ustilago maydis* SG200, SG200Δerc1, SG200Δerc1/UhErc1, SG200Δafu2-3 and SG200Δ3xafu/Erc1 on Early Golden Bantam (EGB) maize cultivar at 12 days post inoculation (dpi). Disease rates are given as a percentage of the total number of infected plants. Two biological replicates were performed for disease assay. n: number of infected maize seedlings.

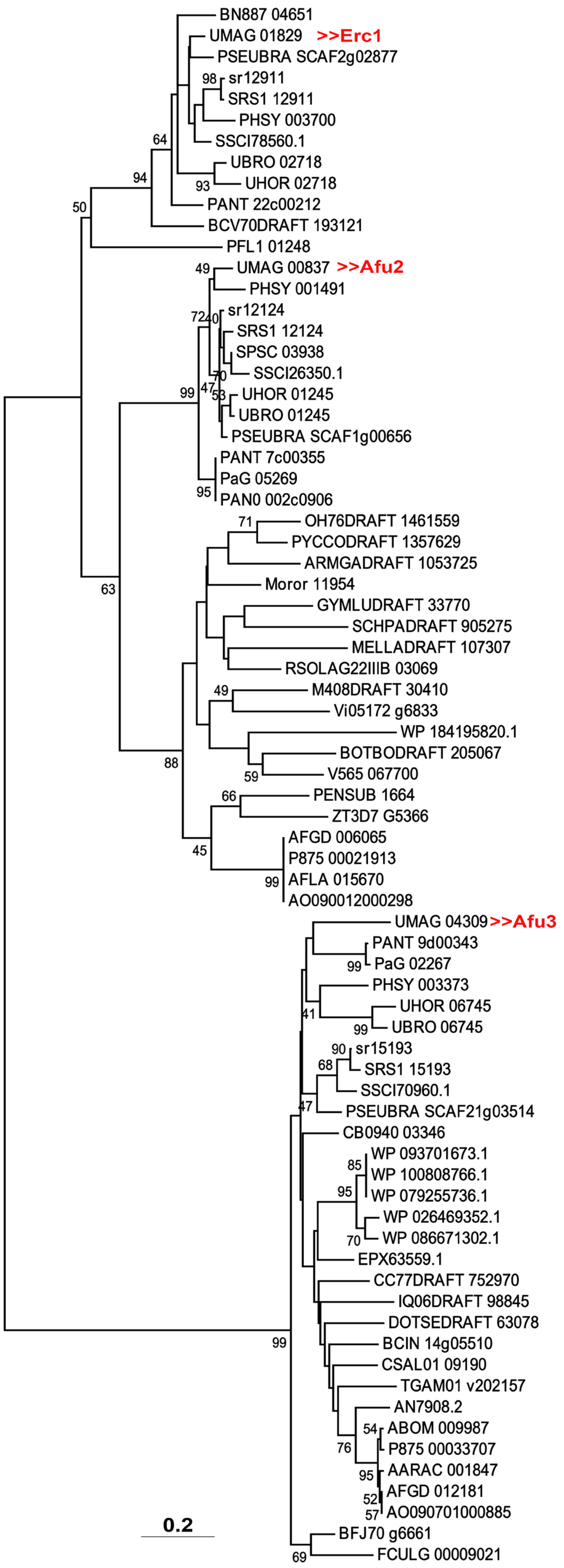

BN887 04651
UMAG 01829 >>Erc1
PSEUBRA SCAF2g02877
sr12911
SRS1 12911
PHSY 003700
SSCI78560.1
UBRO 02718
UHOR 02718
PANT 22c00212
BCV70DRAFT 193121
PFL1 01248
UMAG 00837 >>Afu2
PHSY 001491
sr12124
SRS1 12124
SPSC 03938
SSCI26350.1
UHOR 01245
UBRO 01245
PSEUBRA SCAF1g00656
PANT 7c00355
PaG 05269
PAN0 002c0906
OH76DRAFT 1461559
PYCCODRAFT 1357629
ARMGADRAFT 1053725
Moror 11954
GYMLUDRAFT 33770
SCHPADRAFT 905275
MELLADRAFT 107307
RSOLAG22IIIB 03069
M408DRAFT 30410
Vi05172 g6833
WP 184195820.1
BOTBODRAFT 205067
V565 067700
PENSUB 1664
ZT3D7 G5366
AFGD 006065
P875 00021913
AFLA 015670
AO090012000298
UMAG 04309>>Afu3
PANT 9d00343
PaG 02267
PHSY 003373
UHOR 06745
UBRO 06745
sr15193
SRS1 15193
SSCI70960.1
PSEUBRA SCAF21g03514
CB0940 03346
WP 093701673.1
WP 100808766.1
WP 079255736.1
WP 026469352.1
WP 086671302.1
EPX63559.1
CC77DRAFT 752970
IQ06DRAFT 98845
DOTSEDRAFT 63078
BCIN 14g05510
CSAL01 09190
TGAM01 v202157
AN7908.2
ABOM 009987
P875 00033707
AARAC 001847
AFGD 012181
AO090701000885
BFJ70 g6661
FCULG 00009021
0.2

**Fig. S3.** Phylogenetic tree analyses of GH51 proteins. A minimum evolution tree was constructed by using an alignment of the full-length amino acid sequence of Erc1 homologs obtained from the NCBI database for different microorganisms. The minimum evolution tree was constructed by using the software Mega7 by using the minimum evolution algorithm performing 1000 bootstraps. PHSY_003700: *Pseudozyma hubeiensis* SY62; SRS1_12911: *Sporisorium reilianum* f. sp. *reilianum*; sr12911: *Sporisorium reilianum* SRZ2; SSCI78560.1: *Sporisorium scitamineum*; PANT_22c00212: *Moesziomyces antarcticus* T-34; PSEUBRA_SCAF2g02877: *Kalmanozyma brasiliensis* GHG001; UBRO_02718: *Ustilago bromivora*; UHOR_02718: *Ustilago hordei*; BN887_04651: *Melanopsichium pennsylvanicum*; BCV70DRAFT_193121: *Testicularia cyperi*; PFL1_01248: *Anthracocystis flocculosa* PF-1; PHSY_001491: *Pseudozyma hubeiensis* SY62; PaG_05269: *Moesziomyces aphidis* DSM 70725; Moror_11954: *Moniliophthora roreri* MCA 2997; PYCCODRAFT_1357629: *Trametes coccinea* BRFM310; BOTBODRAFT_205067: *Botryobasidium botryosum* FD-172 SS1; GYMLUDRAFT_33770: *Gymnopus luxurians* FD-317 M1; RSOLAG22IIIB_03069: *Rhizoctonia solani*; ARMGADRAFT_1053725: *Armillaria gallica*; SCHPADRAFT_905275: *Schizopora paradoxa*; AFLA_015670: *Aspergillus flavus*; AFGD_006065: *Aspergillus flavus*; AO090012000298: *Aspergillus oryzae*; P875_00021913: *Aspergillus parasiticus* SU-1; M408DRAFT_30410: *Serendipita vermifera* MAFF 305830; Vi05172_g6833: *Venturia inaequalis*; OH76DRAFT_1461559: *Polyporus brumalis*; PENSUB_1664: *Penicillium subrubescens*; ZT3D7_G5366: *Zymoseptoria tritici* ST99CH_3D7; V565_067700: *Rhizoctonia solani* 123E; PSEUBRA_SCAF1g00656: *Kalmanozyma brasiliensis* GHG001; PAN0_002c0906: *Moesziomyces antarcticus*; SRS1_12124: *Sporisorium reilianum* f. sp. *reilianum*; sr12124: *Sporisorium reilianum* SRZ2; PANT_7c00355: *Moesziomyces antarcticus* T-34; UHOR_01245: *Ustilago hordei*; SPSC_03938: *Sporisorium scitamineum*; UBRO_01245: *Ustilago bromivora*; SSCI26350.1: *Sporisorium scitamineum*; MELLADRAFT_107307: *Melampsora larici-populina* 98AG31; SSCI70960.1: *Sporisorium scitamineum*; sr15193: *Sporisorium reilianum* SRZ2; SRS1_15193: *Sporisorium reilianum* f. sp. *reilianum*; PANT_9d00343: *Moesziomyces antarcticus* T-34; PaG_02267: *Moesziomyces aphidis* DSM 70725; PSEUBRA_SCAF21g03514: *Kalmanozyma brasiliensis* GHG001; UBRO_06745: *Ustilago bromivora*; PHSY_003373: *Pseudozyma hubeiensis* SY62; UHOR_06745: *Ustilago hordei*; WP_086671302.1: *Amycolatopsis pretoriensis*; WP_100808766.1: *Streptomyces* sp. M56; WP_093701673.1: *Streptomyces* sp. MnatMP-M27; WP_079255736.1: *Streptomyces* autolyticus; WP_026469352.1: *Amycolatopsis balhimycina*; EPX63559.1: *Cystobacter fuscus* DSM 2262; WP_184195820.1: *Armatimonas rosea*; AARAC_001847: *Aspergillus arachidicola*; ABOM_009987: *Aspergillus bombycis*; AO090701000885: *Aspergillus oryzae* RIB40; BCIN_14g05510: *Botrytis cinerea* B05.10; P875_00033707: *Aspergillus parasiticus* SU-1; AFGD_012181: *Aspergillus flavus*; AN7908.2: *Aspergillus nidulans* FGSC A4; TGAM01_v202157: *Trichoderma gamsii*; CSAL01_09190: *Colletotrichum salicis*; BFJ70_g6661: *Fusarium oxysporum*; FCULG_00009021: *Fusarium culmorum*; DOTSEDRAFT_63078: *Dothistroma septosporum* NZE10; CB0940_03346: *Cercospora beticola*; IQ06DRAFT_98845: *Stagonospora* sp. SRC1lsM3a; CC77DRAFT_752970: *Alternaria alternata*

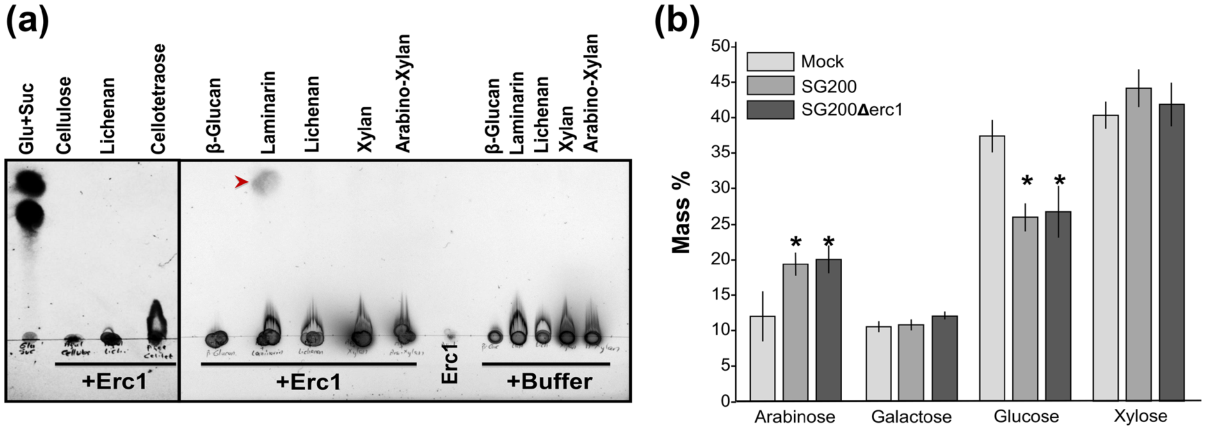


**Fig. S4. (a) Thin-layer chromatography (TLC) assay** to show enzymatic activity of Erc1 on different polysaccharides, including cellulose, lichenan, cellotetraose, β-glucan, laminarin, xylan and arabinoxylan. A mixture of n-propanol:ethanol:water (7:2:1/v:v:v) was used as a mobile phase. Carbohydrates and their hydrolysis products were visualized by spraying the TLC plate with detection solution and subsequent drying at 100°C for approximately 15 min. Red arrow head indicates hydrolysis product of laminarin. Glucose + sucrose mix was used as reference. **(b) Monosaccharide composition of maize cell walls.** Plant cell walls were isolated from mock, SG200 and SG200Δerc1-treated EGB maize leaves. The isolated plant cell walls were analyzed for their arabinose, galactose, glucose and xylose content. Asterisks above bars indicate significant differences ($p<0.05$, student t-test).

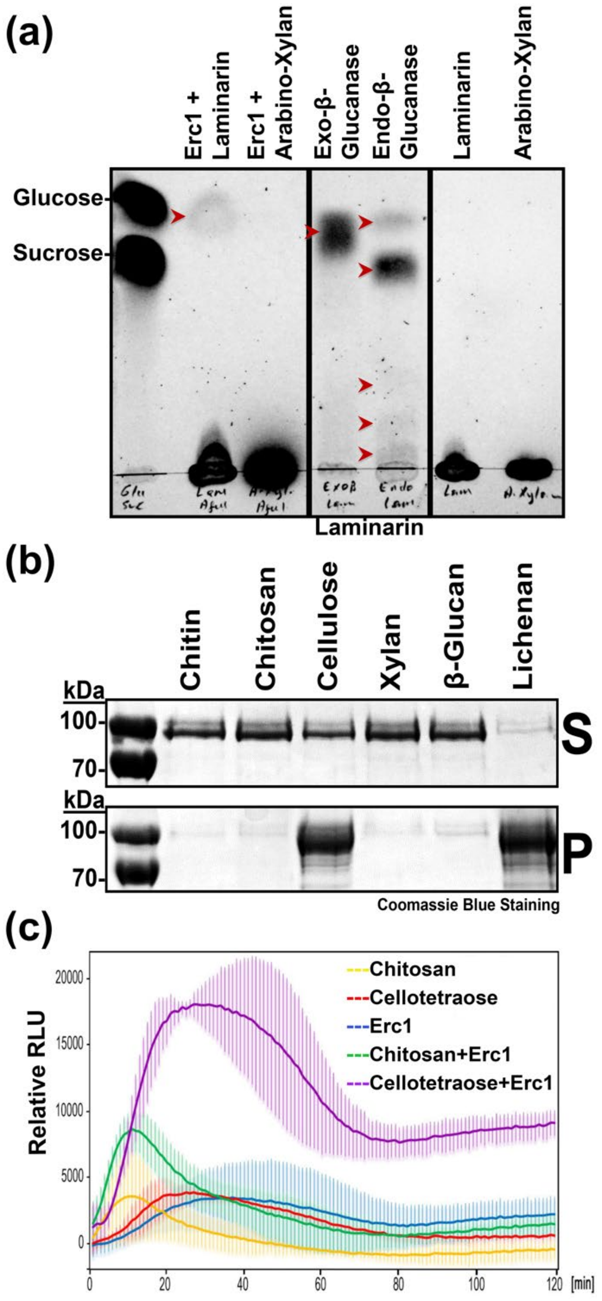


**Fig. S5. (a)** Thin-layer chromatography (TLC) assay was performed to demonstrate exo-β-1,3-glucanase activity of Erc1 on laminarin. Glucose + sucrose mix was used as reference. A mixture of n-propanol:ethanol:water (7:2:1/v:v:v) was used as a mobile phase. Laminarin and its hydrolysis products were visualized by spraying the TLC plate with detection solution and subsequent drying at 100°C for approximately 15 min. Red arrow heads indicate hydrolysis products of laminarin. Commercial exo- and endo-β-1,3-glucanases were used as controls. **(b)** Carbohydrate binding assay for recombinant Erc1 protein. Erc1 protein was incubated with insoluble chitin, chitosan, cellulose, xylan, β-glucan and lichenan. Subsequently, supernatant and pellet phases were analysed for presence of Erc1 protein via SDS-PAGE followed by coomassie blue staining. S: supernatant, P: pellet **(c)** To check whether Erc1 sequesters cellotetraose in order to prevent its recognition as a DAMP, a ROS-burst assay was performed with barley leaf disks incubated with cellotetraose, Erc1 and cellotetraose incubated with Erc1 recombinant protein. Chitosan was used as a negative control. Relative luminescence units (RLU) indicate ROS burst activity of treated barley leaf discs. The RLU are normalized with the buffer control. The error bars show the standard deviation of the three replicates.

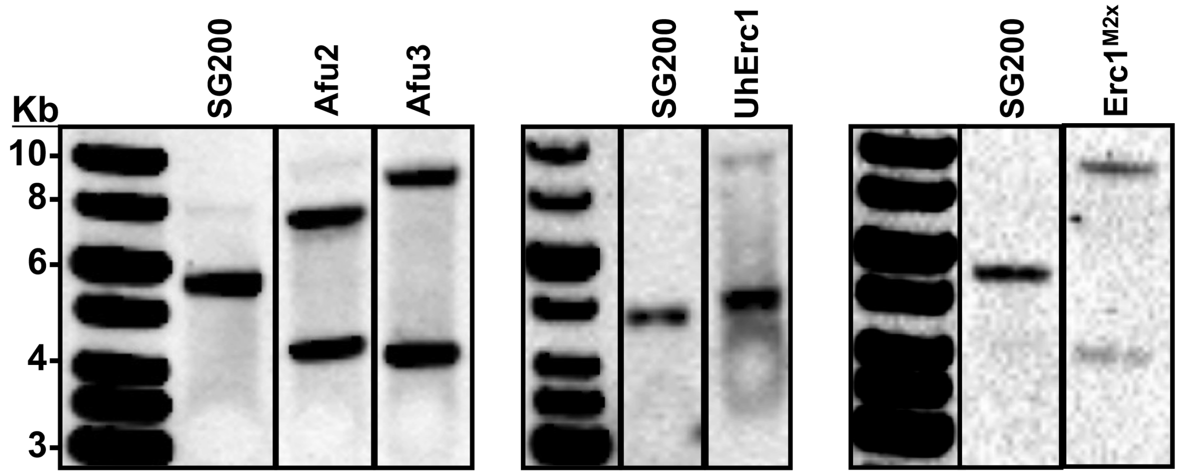


| Cbx integration | Afu2 (*Bam*HI) | Afu3 (*Bam*HI) | UhErc1 (*Sac*I) | UhErc1$^{M2x}$ (*Bam*HI) |
|---|---|---|---|---|
| Wild type: | 5662 bp | 5662 bp | 4756 bp | 5662 bp |
| Single integration: | 4065-6534 bp | 4061-8134 bp | 3921-6949 bp | 4073-9352 bp |
| Multiple integration: | 4065-4937-6534 bp | 4061-6533-8134bp | 3921-6114-6949 bp | 4073-7763-9352 bp |

**Fig. S6. Southern Blot analysis to confirm a single insertion event in different SG200Δerc1 complementation strains**. gDNA of both SG200 and complementation strains were digested with *Bam*HI or *Sac*I restriction enzyme. Cbx gene was used as a probe to detect single insertion event. Expected band sizes for wild-type, single and multiple integration events are depicted below the figure.

**Supporting Table**

**TableS1. Primer and construct list.**

**Supporting Information: Materials and Methods S1:** The experiments performed in this manuscript have been explained in more detail.

**Growth conditions for fungal and bacterial cultures**

The *Escherichia coli* DH5α strain was grown in dYT-medium (1.6% w $v^{-1}$ peptone, 1% w $v^{-1}$ yeast extract and 0.5% w $v^{-1}$ NaCl) with appropriate antibiotics at 37°C with 200 rpm shaking (for liquid cultures). *Ustilago maydis* SG200 and *U. hordei* DS199 solopathogenic strains were incubated in $YEPS_{light}$ (0.4% w $v^{-1}$ yeast extract, 0.4% w $v^{-1}$ peptone, and 2% w $v^{-1}$ sucrose) liquid medium at 28°C and 20°C with 200 rpm shaking, respectively (Ökmen *et al.*, 2021). Growth of *U. maydis* and *U. hordei* cultures on plates was carried out on potato dextrose agar with appropriate antibiotics (concentrations: 200 µg $ml^{-1}$ hygromycin or 2 µg $ml^{-1}$ carboxin). The *Pichia pastoris* KM71H-OCH strain was used for recombinant protein expression. YPD medium supplemented with 100 µg $ml^{-1}$ zeocin was used for the initial growth of *P. pastoris* strains at 28°C and 200 rpm shaking (for liquid cultures). *Zea mays* L. Early Golden Bantam (EGB) maize (Olds Seeds, Madison, WI, USA) and *Hordeum vulgare* Golden Promise (GP) barley cultivars were grown for infection assays.

**Nucleic acids methods**

Plasmid DNA from bacterial cells was isolated using the QIAprep Spin Miniprep Kit (Qiagen; Hilden, Germany) according to the manufacturer's information. The genomic DNA isolation from *U. maydis* and *U. hordei* was performed according to the protocol described by Schultz *et al.* (1990). The polymerase chain reaction (PCR) utilizing Phusion© polymerase enzyme (Thermo Fisher Scientific;Darmstadt, Germany) was used to amplify specific DNA fragments by using gene specific primer pairs depicted in **Table S1**.

Total RNA isolation was performed with crushed infected leaf material (at 4 dpi) using the TRIzol® extraction method (Invitrogen; Karlsruhe, Germany) according to the manufacturer's instructions. Subsequently, the Turbo DNA-Free™ Kit (Ambion/Applied Biosystems; Darmstadt, Germany) was used to remove any genomic DNA contamination. cDNA synthesis was performed with 1 µg of total isolated RNA by using the First strand cDNA synthesis Kit (Thermo Fisher Scientific; Darmstadt, Germany). RT-qPCR analysis for *PR* gene expression was performed by using SYBR® Green Supermix (BioRad; Munich, Germany). The reaction was performed in a Bio-Rad iCycler system using the following program: 2 min at 95°C followed by 45 cycles of 30 s at 95°C, 30 s at 61°C and 30 s 72°C. The expression levels of maize *PR* genes were calculated relative to the *GAPDH* gene of maize (*NM001111943*). Results of at least three biological RT-qPCR replicates were analyzed using the $2^{-\Delta Ct}$ method (Livak & Schmittgen, 2001). The primers used for RT-qPCR are listed in **Table S1**.

***Ustilago maydis* virulence assays**

The *U. maydis* virulence assays on the EGB maize cultivar were performed as described in Redkar and Doehlemann (2016). Disease symptoms for *U. maydis* were scored at 12 dpi using the disease rating scheme developed previously (Kämper *et al.*, 2006). For statistical analysis for *U. maydis* virulence assays, the disease index was calculated as follows: The number of plants sorted into categories 'chlorosis', 'small tumor', 'normal tumor', 'big tumor' and 'heavy tumor' were multiplied by 1, 3, 6, 9 and 12, respectively. All calculated numbers for each strain were summed and then divided by the total number of infected plants. The *U. hordei* virulence assays on the GP barley cultivar were performed as described in Ökmen *et al.*, (2018). Infected barley leaves were collected at 8 dpi for gDNA isolation and followed by qPCR to quantify the fungal biomass in mutant and DS199 strains. All virulence assays were performed in three independent biological replicates. A student t-test was performed to calculate significant differences in disease indices between mutant and solopathogenic strains.

**Heterologous protein production in *Pichia pastoris***

The *Pichia pastoris* KM71H-OCH protein expression system was used to produce N-terminally His and C-terminally Myc-His tagged *U. maydis* Erc1, Erc1$^{M1x}$, and Erc1$^{M2x}$ recombinant proteins. All genes for each respective protein were cloned into the pGAPZαA vector (Invitrogen; Carlsbad, USA) under the control of a constitutive promotor with an α-factor signal peptide for secretion. Protein expression was performed by growing *Pichia* in 1 L buffered (100 mM sodium phosphate buffer, pH 6.0) YPD medium at 28°C for 48 hours with 200 rpm shaking (pGAPZαA, B, & C *Pichia pastoris* Expression Vectors, Invitrogen; Carlsbad, USA). Recombinant protein purification was performed with a Ni-NTA-matrix (Ni-Sepharose™ 6 Fast-Flow, GE-Healthcare; Freiburg, Germany). After protein purification, each protein sample was applied to the NAP-25 column to exchange buffer with 20 mM potassium phosphate buffer pH 6.0. The proteins were stored at -20°C for further experiments. Western blot analysis was performed as described in Mueller *et al.* (2013) using anti-His and anti-Myc antibodies.

**Enzyme activity assay and activity-based protein profiling**

To determine the enzymatic activity of the purified Erc1 protein, 4-nitrophenyl α-L-arabinofuranoside (4NPA) (Sigma-Aldrich, Steinheim) was diluted in 0.1 M sodium acetate buffer (pH 4) to concentrations of 1 mM and 5 mM. The 4NPA substrate was incubated with 1.34 and 9 µM Erc1 and heat inactivated recombinant protein at 40°C. After 10 min incubation, the reaction was stopped by adding 150 µl 2% trisodium phosphate buffer (pH: 12). The absorption was measured at 400 nm. The commercially available arabinofuranosidase (AFASE) from *Aspergillus niger* (Megazyme, Ireland) was used as a positive control.

For activity-based protein profiling (ABPP) assay, 15 µl (80 µg ml$^{-1}$ stock) of Erc1, Erc1$^{M2x}$ and AFASE recombinant proteins were incubated in 300 mM sodium acetate buffer pH:6.0 for 30 minutes with and without 50 µM DL69 α-L-arabinofuranosidase specific inhibitor (McGregor *et al.*, 2020). Subsequently, 5 µM α-L-arabinofuranosidase specific ME868 probe was added to the reaction mixture and incubated for 2 hours at room temperature at dark. The reaction was stopped with 6X SDS-loading dye and the probe was detected by scanning the in-gel fluorescence with Cy5 filter (Ex. 650 nm, Em. 670 nm). Protein of loaded samples were visualized via Sypro Ruby (Ex. 450 nm, Em. 610 nm).

**Thin-layer chromatography (TLC) assay**

To detect any carbohydrate hydrolysing activity of Erc1 a thin-layer chromatography assay (TLC) was performed with different polysaccharides incubated with purified Erc1 recombinant protein as described in Ökmen *et al*. (2019). Erc1 (50 µl, 500 µg $ml^{-1}$) recombinant protein was incubated with 50 µl of different carbohydrate solutions (5 mg $ml^{-1}$) including β-glucan (from barley, Megazyme, lot: 90803b), laminarin (from *Laminaria digitate*, Sigma, L9634), lichenan (from moss, Megazyme, lot: 80402), xylan (from beechwood, Megazyme, lot: 171002) and arabino-Xylan (from wheat, Megazyme, lot: 120601b) overnight at 42˚C. For another TLC assay, 25 µL Erc1 (500 µg $ml^{-1}$), 25 µl $Erc1^{M1x}$ (500 µg $ml^{-1}$) and 36 µL $Erc1^{M2x}$ (350 µg $ml^{-1}$) recombinant protein was incubated with 5 µl of laminarin (from *Laminaria digitate*, Sigma, L9634) and laminarihexaose (Megazyme, lot: 190606) solutions (5 mg $ml^{-1}$ per each) in 70 µl water overnight at 42˚C. After spinning down the insoluble polysaccharides, 20 µl of each digest was loaded on a TLC Silica gel 60 $F_{254}$ plate (20x20 cm) (Merck, HX85205954). Two µL of 3 mg $m^{-1}$ Glucose + Sucrose mixture was used as reference. Untreated polysaccharide sample and recombinant protein alone samples were used as negative controls. n-propanol:ethanol:water (7:2:1) (v:v:v) was used as running solvent for the TLC assay. Carbohydrates were visualized by spraying a staining solution **(**45 mg naphthol in 4.8 ml sulfuric acid, 37.2 ml ethanol, 3 ml water) onto the dried TLC plate and subsequent 5-10 min incubation of sprayed TLC plate at 100°C.

**WGA-AF488/Propidium iodide staining of infected plant tissues**

The WGA-AF488 (Molecular Probes, Karlsruhe, Germany) and propidium iodide (Sigma-Aldrich) cell staining was performed according to Ökmen *et al*. (2018). WGA-AF488 stains fungal cell walls (green), while the propidium iodide stains plant cell walls (red). Briefly, infected leaf material was bleached in pure ethanol and subsequently boiled for 1-2 hours in 10% KOH at 85°C. The pH of the leaf samples was neutralized using 1xPBS buffer (pH: 7.4) with several washing steps. The WGA-AF488/PI staining solution (1 µg $ml^{-1}$ propidium iodide, 10 µg $ml^{-1}$ WGA-AF 488; 0.02% Tween 20 in PBS pH 7.4) was vacuum infiltrated into leaf samples three times with a desiccator for 5 min at 250 mbar. WGA-AF488: excitation at 488 nm; detection at 500-540 nm. PI: excitation at 561 nm; detection at 580–630 nm.

**Transmission electron microscopy and immunogold labelling**

**High pressure freezing, freeze substitution and resin embedding**

For TEM observations, 2 mm leaf discs from maize leaves infected with *U. maydis* expressing Erc1-HA and GFP-HA under the control of the native Erc1 promoter and signal peptide, were excised from 1 cm below infected area using a biopsy punch and processed by means of high pressure freezing (HPF) following the procedure described in Micali *et al*. (2011).

Freeze substitution (FS) was performed in 0.5% uranyl acetate in acetone (w/v) in a Leica EM AFS2 freeze substitution device (Leica Microsystems GmbH) over 7 days with temperatures ranging from -85°C (90 h) over -60°C (24 h) and -30°C (24 h) to -20°C (12 h). At the end of the FS run, samples were rinsed in acetone and gradually transferred into ethanol into a Leica EM AFS2 at -20°C and then carefully removed from their HPF carriers in ice-cold ethanol with the help of a stereomicroscope. Subsequent infiltration in medium-grade LR White resin (Plano GmbH) was gradually performed over 7 days in a freezer at -20°C with the help of a laboratory rocker; LR White polymerization with UV light in the Leica EM AFS2 was achieved for 24 h at -20°C and 24 h at 0°C.

**Sectioning, immunogold labelling and transmission electron microscopy**

Ultrathin (70-90 nm) sections were collected on nickel slot grids as described by Moran and Rowley (1987). For immunogold labelling, sections were blocked for 30 min in a 1:30 dilution of goat normal serum in TRIS buffer (20 mM TRIS, 15 mM NaN3, 225 mM NaCl, pH 6.9) supplemented with 1% (w/v) BSA (Sigma-Aldrich A3294) and 1% (w/v) fish gelatin (FG; Sigma-Aldrich G7765). After three washes for 10 min in TRIS-BSA-FG, sections were incubated in a 1:500 dilution of monoclonal mouse anti-HA antibody (Sigma Aldrich H-9658) for 1 h at room temperature (slow orbital shaking). After washing in TRIS-BSA-FG (4× 10 min), sections were incubated in a 1:20 dilution of secondary goat anti-mouse antibody conjugated to 10 nm colloidal gold particles (British Biocell International, Cardiff, UK) for 1 h (slow orbital shaking). Finally, sections were rinsed in TRIS-BSA-FG (4x 5 min) followed by a stream of sterile-filtered, distilled water for 3 min. After drying at room temperature, sections were examined without further staining in a Hitachi H-7650 TEM (Hitachi High-

Technologies Europe GmbH, Krefeld, Germany) operating at 100 kV fitted with an AMT XR41-M digital camera (Advanced Microscopy Techniques, Danvers, USA).

**Bioinformatics methods**

The SignalP 5 software program was used to predict an N-terminal secretion signal peptide within a protein sequence (http://www.cbs.dtu.dk/services/SignalP/). For protein domain prediction the website Pfam (http://pfam.xfam.org/) was used. A minimum evolution tree was constructed by using an alignment of the full-length amino acid sequence of Erc1 homologs obtained from the NCBI database for different microorganisms. The minimum evolution tree was constructed by using the software Mega7 (http://www.megasoftware.net/mega.php) with a minimum evolution algorithm performing 1000 bootstraps.
